# Durable immunity to SARS-CoV-2 in both lower and upper airways achieved with a gorilla adenovirus (GRAd) S-2P vaccine in non-human primates

**DOI:** 10.1101/2023.11.22.567930

**Authors:** Juan I. Moliva, Shayne F. Andrew, Barbara J. Flynn, Danielle A. Wagner, Kathryn E. Foulds, Matthew Gagne, Dillon R. Flebbe, Evan Lamb, Samantha Provost, Josue Marquez, Anna Mychalowych, Cynthia G. Lorag, Christopher Cole Honeycutt, Matthew R. Burnett, Lauren McCormick, Amy R. Henry, Sucheta Godbole, Meredith E. Davis-Gardner, Mahnaz Minai, Kevin W. Bock, Bianca M. Nagata, John-Paul M. Todd, Elizabeth McCarthy, Alan Dodson, Katelyn Kouneski, Anthony Cook, Laurent Pessaint, Alex Van Ry, Daniel Valentin, Steve Young, Yoav Littman, Adrianus C. M. Boon, Mehul S. Suthar, Mark G. Lewis, Hanne Andersen, Derron A. Alves, Ruth Woodward, Adriano Leuzzi, Alessandra Vitelli, Stefano Colloca, Antonella Folgori, Angelo Raggiolli, Stefania Capone, Martha C. Nason, Daniel C. Douek, Mario Roederer, Robert A. Seder, Nancy J. Sullivan

## Abstract

SARS-CoV-2 continues to pose a global threat, and current vaccines, while effective against severe illness, fall short in preventing transmission. To address this challenge, there’s a need for vaccines that induce mucosal immunity and can rapidly control the virus. In this study, we demonstrate that a single immunization with a novel gorilla adenovirus-based vaccine (GRAd) carrying the pre-fusion stabilized Spike protein (S-2P) in non-human primates provided protective immunity for over one year against the BA.5 variant of SARS-CoV-2. A prime-boost regimen using GRAd followed by adjuvanted S-2P (GRAd+S-2P) accelerated viral clearance in both the lower and upper airways. GRAd delivered via aerosol (GRAd(AE)+S-2P) modestly improved protection compared to its matched intramuscular regimen, but showed dramatically superior boosting by mRNA and, importantly, total virus clearance in the upper airway by day 4 post infection. GrAd vaccination regimens elicited robust and durable systemic and mucosal antibody responses to multiple SARS-CoV-2 variants, but only GRAd(AE)+S-2P generated long-lasting T cell responses in the lung. This research underscores the flexibility of the GRAd vaccine platform to provide durable immunity against SARS-CoV-2 in both the lower and upper airways.

## Introduction

Severe acute respiratory syndrome coronavirus-2 (SARS-CoV-2) is an infectious respiratory human coronavirus and the causative agent of COVID-19. At the time of writing, COVID-19 has been responsible for more than 676 million cases and 6.8 million deaths worldwide.^1^ In response to its rapid spread, vaccine development was substantially accelerated, resulting in the evaluation and approval of numerous novel vaccine technologies, namely the mRNA encapsulated (*e.g*., mRNA-1273 and BNT162b2) and viral-vectored (*e.g*., ChAdOx nCoV-19/AZD1222 and Ad26.COV2.S) platforms.^2^ Initially, and for much of the pandemic, the vaccines approved for human use were based on the ancestral Wuhan-Hu-1/USA-WA1/2020 (WA-1) isolate and proved remarkably effective against severe disease.^3^ However, the high transmissibility of SARS-CoV-2 facilitated the rise of mutations, deletions and insertions in the Spike (S) protein, the S receptor-binding domain (RBD) and the S N-terminal domain (NTD) leading to the emergence of numerous subvariants with progressive resistance to vaccine-elicited immunity, diminishing the durable efficacy of the vaccines and increasing transmissibility.^4–7^

One significant limitation of most SARS-CoV-2 vaccines, including all mRNA-based vaccines approved for human use to date, is that they do not directly stimulate immune responses in the mucosal surfaces of the upper and lower airway, the major site of SARS-CoV-2 entry, replication, pathology, and shedding.^8^ Despite this, intramuscular mRNA-based vaccines induce robust humoral and cellular immunity and are effective against severe disease caused by SARS-CoV-2.^9^ However, the ability of intramuscularly delivered mRNA-based vaccines to prevent transmission is limited,^10,11^ and efficacy against severe disease wanes,^12^ possibly due to the failure of properly priming the respiratory mucosa. Since the respiratory tract serves as the first line of defense against SARS-CoV-2, robust and durable immunity in the mucosa could provide systemic durable immunity and prevent initial infection and subsequent transmission as tissue-resident memory T and B cells are likely among the early responders.^13^ Indeed, pre-clinical studies have demonstrated that mucosal immunity can protect against SARS-CoV-2.^14–16^ Therefore, it is important to evaluate additional vaccine approaches and candidates that can prime or boost mucosal immunity.

GRAd32 is a novel replication-defective simian adenoviral vector that was isolated from a captive gorilla.^17^ Simian adenoviruses do not cause disease in humans, and consequently, have low or no seroprevalence in the human population making them suitable vaccine candidates.^18^ Additionally, simian-based adenoviral vaccine vectors have been proven safe in humans, generate potent and durable immune responses, and have been shown to stimulate humoral and cellular responses across a plethora of antigens (*e.g.*, Ebola, malaria, hepatitis C, human immunodeficiency virus, and respiratory syncytial virus, SARS-CoV-2).^19–22^ When combined with other vaccine platforms (*e.g.,* mRNA or subunit protein), called heterologous immunization strategies, the inclusion of a viral-vectored vaccine can confer superior durability and efficacy compared to homologous vaccination.^23–26^ In this study, we used the GRAd32 vector backbone encoding the WA-1 SARS-CoV-2 pre-fusion stabilized S (henceforth abbreviated GRAd in this manuscript) to assess the durability of immune responses in a combination of prime and prime and boost homologous and heterologous immunization strategies to evaluate the protective efficacy to SARS-CoV-2.

Using non-human primates (NHP), a model that has faithfully predicted protective efficacy of SARS-CoV-2 vaccines in humans,^27,28^ herein we demonstrate efficacy of the GRAd vaccine platform encoding WA-1 against mismatch challenge with BA.5. Groups of eight NHP were primed with intramuscular (IM) GRAd, primed and boosted with a combination of IM GRAd and adjuvanted S-2P (GRAd+S-2P), primed and boosted with IM adjuvanted S-2P (S-2P+S-2P), or primed with aerosolized (AE) GRAd and boosted with IM adjuvanted S-2P (GRAd(AE)+S-2P). The NHP were followed for 48 weeks, at which point four of the eight NHP were boosted with mRNA BNT162b2 (S-2P encoded in mRNA). At week 64, all NHP were challenged with SARS-CoV-2 BA.5. For the duration of the 64 weeks, we collected blood, bronchoalveolar lavage (BAL) and nasal washes to analyze systemic and mucosal antibody kinetics and B and T cell responses. Following BA.5 challenge, viral replication in the lung and nose was quantified to assess the efficacy of these vaccine regimens. We found that GRAd in any combination conferred year-long protection in the lower airway, but heterologous GRAd vaccine regimens were superior. Priming with AE GRAd provided an advantage to IM GRAd in control of BA.5 in the upper airway.

## Results

### GRAd confers durable protection against BA.5 in the lower airway

To quantify the durable protective effect of GRAd or adjuvanted protein subunit-based immunogens against SARS-CoV-2, Indian-origin rhesus NHP were stratified into groups of eight and immunized according to the experimental schema (**Fig. S1A**). At week 48, all groups (except control) were further subdivided, and four of the eight NHP were boosted with BNT162b2 (henceforth abbreviated mRNA in this manuscript). At week 64, each NHP was challenged with a dose of 8 × 10^5^ PFU of sequence-confirmed Omicron sub-lineage BA.5 (**Fig. S2A-D**). BAL was collected on days 2, 4, and 8 (**Fig. S1B**). SARS-CoV-2 subgenomic RNA (sgRNA) in BAL was quantified to assess the extent of viral burden in the lower airway. Specifically, we relied primarily on the sgRNA encoding for the N gene (sgRNA_N) as it is the most abundant transcript produced due to the viral discontinuous transcription process, and thus provides the most sensitive way to detect SARS-CoV-2.^29^

Two days following challenge, control NHP had a geometric mean copy number of 168261 sgRNA_N per mL (**Fig. 1A**), whereas GRAd, S-2P+S-2P, GRAd+S-2P and GRAd(AE)+S-2P had 3091, 4690, 138 and 81, respectively, representing statistically significant decreases except in the S-2P+S2P group (**Fig. 1B**). By day 4, the copy number in the control NHP was 102535, and significantly lower in all immunized groups except for S-2P+S-2P (GRAd −144; S-2P+S-2P - 1111; GRAd+S-2P - 135; GRAd(AE)+S-2P - 77). By day 8, copy number in the control NHP was 2244, below 100 in the GRAd and S-2P+S-2P groups, and below 50 (limit of detection) in the GRAd+S-2P and GRAd(AE)+S-2P groups. Altogether, the data suggest year-long and rapid protection can be achieved in the lower airway with a prime and boost strategy that includes one dose of GRAd.

**Figure 1.**
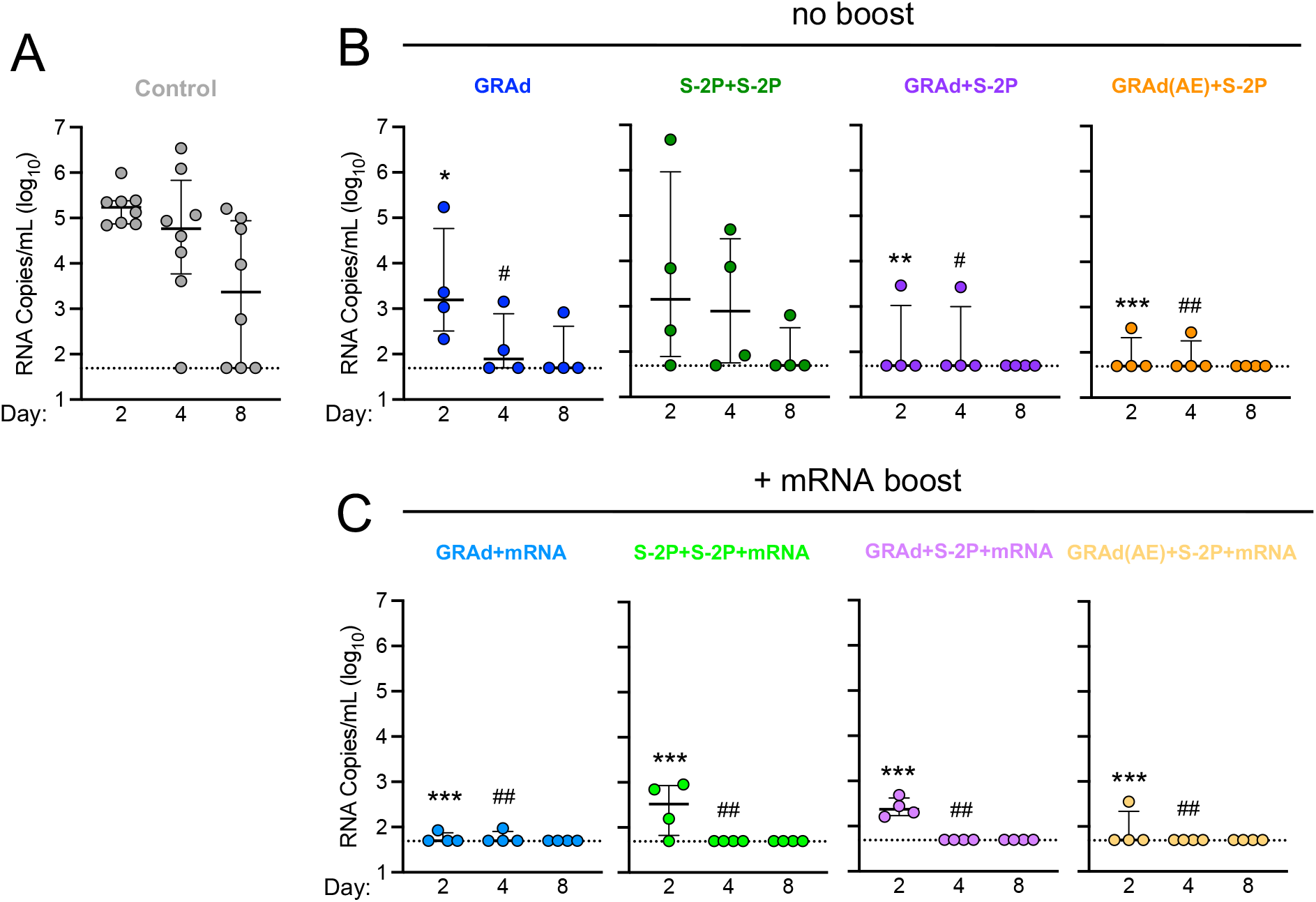
GRAd confers durable protection against BA.5 in the lower airway. (A–C) BAL was collected at days 2, 4 and 8 following challenge with 8 × 10^5^ PFU BA.5. (A) BA.5 sgRNA_N copy numbers per mL of BAL in control NHP. (B) BA.5 sgRNA_N copy numbers per mL of BAL in GRAd, S-2P+S-2P, GRAd+S-2P and GRAd(AE)-S-2P NHP. (C) BA.5 sgRNA_N copy numbers per mL of BAL in GRAd, S-2P+S-2P, GRAd+S-2P and GRAd(AE)-S-2P NHP boosted with mRNA at week 48. Circles (A–C) indicate individual NHP. Error bars represent interquartile range with the median denoted by a horizontal line. Assay limit of detection indicated by a dotted horizontal line. Statistical analysis shown for corresponding timepoints between control and test group (*e.g*., ‘*’ symbols denote comparisons at day 2, ‘^#^’ symbols denote comparison at day 4). *,^#^ *p* <0.05, **,^##^ *p* <0.01, *** *p* <0.001. Eight control NHP and 4 immunized NHP per cohort. See also Figure S1 for experimental schema, Figure S2 for BA.5 titration in NHP, Figure S3 for viral load and Figure S4 for lung pathology.

We next asked whether the mRNA boost amplified the observed protection. The mRNA boost had its greatest impact on the GRAd and S-2P+S-2P groups, reducing the day 2 geometric mean sgRNA copy number from 3091 to 57 and 4690 to 264, respectively, while the sgRNA_N copy number remained approximately equal in the GRAd+S-2P and GRAd(AE)+S-2P groups at day 2. By day 4, all mRNA-boosted NHP had a nearly undetectable virus in the lower airway, and no sgRNA_N was detected in any group at day 8 (**Fig. 1C**). This data suggests a limited benefit of additional mRNA boost if the priming immunization is adequate.

In addition to measuring the sgRNA_N, we quantified the amount of culturable virus in the BAL using a tissue culture infectious dose assay (TCID_50_). While BA.5 was recovered from 7/8, 6/8, and 2/8 control NHP at days 2, 4 and 8, respectively (**Fig. S3A**), hardly any virus was recovered from the immunized groups, irrespective of whether they received the mRNA boost (**Fig. S3B, C**). Of note, the only group with no recoverable virus at any timepoint in the BAL were the NHP that received GRAd(AE)+S-2P as the primary immunization series.

Finally, to assess pathology caused by SARS-CoV-2 BA.5 to the lung, two NHP from each group were euthanized on days 8 or 9 following challenge, and the extent of viral antigen and inflammation were quantified. SARS-CoV-2 antigen was detected in variable amounts in the lung of 3/4 control NHP and only associated with the alveolar septa lining in areas of interstitial inflammation and thickening. No antigen was detected in the lung of any of the immunized NHP on days 8 or 9, nor in any of the NHP on days 14 or 15 (**Fig. S4A**). Histopathologic lesions were observed in one or more lung lobes in one or more NHP from each group on days 8 or 9 and 14 or 15. Generally, the inflammatory lesion severity in immunized NHP ranged from minimal to moderate, although more severe changes were occasionally observed, while that of control NHP ranged from moderate to severe. Mild to moderate inflammation was characterized by few foci of perivascular inflammation or cuffing without other changes. Increases in severity were marked as foci of perivascular inflammation accompanied by mild alveolar interstitial thickening by mononuclear cells, protein-rich edema accumulation in the alveolar lumina, minimal type II pneumocyte hyperplasia, and mild histiocytic and neutrophilic inflammation to the alveolar lumina. Further increases in severity had a more widespread distribution of the listed lesions. Moderate to marked neutrophilic peribronchial infiltrates were also observed in the highly affected lobes (**Fig. S4B**).

### Aerosol delivery of GRAd rapidly protects against BA.5 in the upper airway

Nasal swabs (NS) were collected from each NHP on days 2, 4 and 8 (**Fig. S1B**) to assess viral burden in the upper airway, the primary source of transmission for SARS-CoV-2.^30^ Two days following challenge, control NHP had geometric mean copy number of 41186 sgRNA_N per swab (**Fig. 2A**), with 1/8 NHP having no detectable virus, whereas GRAd, S-2P+S-2P, GRAd+S-2P and GRAd(AE)+S-2P had 66251, 21952, 7541 and 1151, respectively (**Fig. 2B**). Though not significant, GRAd(AE)+S-2P NHP had a median sgRNA_N copy number 35-fold lower than control NHP on day 2. By day 4, the copy number in the control NHP was 22960 and had decreased substantially, albeit non-significantly, in the GRAd+S2P and GRAd(AE)+S-2P immunized groups (GRAd - 91560; S-2P+S-2P - 32276; GRAd+S-2P - 2738; GRAd(AE)+S-2P 335). Furthermore, on day 4, 2/4 GRAd+S-2P NHP and 3/4 GRAd(AE)+S-2P had undetectable sgRNA_N in the nose. By day 8, copy number in the control NHP was 12820 but was significantly lower in the GRAd+S-2P and GRAd(AE)+S-2P groups (GRAd - 2491; S-2P+S-2P 1918; GRAd+S-2P - 122; GRAd(AE)+S-2P - 71). Together, the data suggest that a prime and boost immunization strategy with GRAd can rapidly reduce the viral load in the nose (**Fig. 2B**).

**Figure 2.**
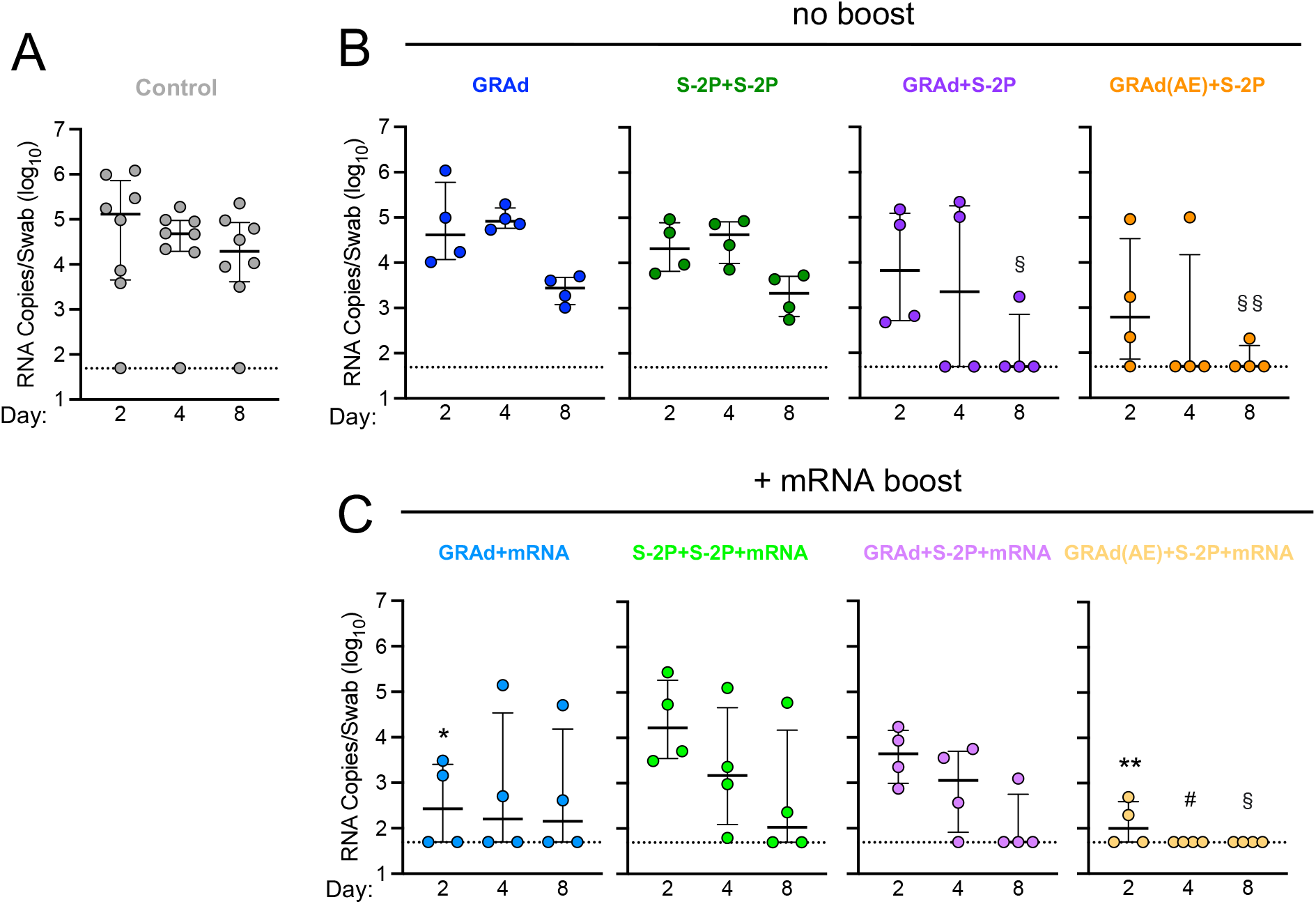
Aerosol delivery of GRAd rapidly protects against BA.5 in the upper airway. (A-C) NS was collected at days 2, 4 and 8 following challenge with 8 × 10^5^ PFU BA.5. (A) BA.5 sgRNA_N copy numbers per swab in control NHP. (B) BA.5 sgRNA_N copy numbers per swab in GRAd, S-2P+S-2P, GRAd+S-2P and GRAd(AE)-S-2P NHP. (C) BA.5 sgRNA_N copy numbers per swab in GRAd, S-2P+S-2P, GRAd+S-2P and GRAd(AE)-S-2P NHP boosted with mRNA at week 48. Circles (A–C) indicate individual NHP. Error bars represent interquartile range with the median denoted by a horizontal line. Assay limit of detection indicated by a dotted horizontal line. Statistical analysis shown for corresponding timepoints between control and test group (*e.g*., ‘*’ symbols denote comparison at day 2, ‘^#^’ symbols denote comparison at day 4, ‘^§^’ symbols denote comparison at day 8). *,^#^,^§^ *p* <0.05, **,^§§^ *p* <0.01. Eight control NHP and 4 immunized NHP per cohort. See also Figure S1 for experimental schema, Figure S2 for BA.5 titration in NHP and Figure S3 for viral load.

Next, we asked whether an mRNA boost could improve the observed protection in the upper airway. Boosting with mRNA significantly decreased the sgRNA_N copy number on day 2 in the GRAd and GRAd(AE)+S-2P groups, reducing the geometric mean copy number from 66251 to 324 and 1151 to 125, respectively. The mRNA boost had no significant impact on the sgRNA_N copy number in the S-2P+S-2P and GRAd+S-2P groups on day 2. By day 4, the GRAd, S-2P+S-2P and the GRAd+S-2P group had detectable virus in the nose, while 4/4 NHP in the GRAd(AE)+S-2P group had cleared the virus. By day 8, BA.5 virus remained detectable in all groups except in the GRAd(AE)+S-2P NHP (**Fig. 2C**). The TCID_50_ assay confirmed these findings (**Fig. S3D-F**). Together, the data suggest that rapid clearance of SARS-CoV-2 from the nasal passage can be achieved with a vaccine regimen that includes AE delivery of GRAd.

### GRAd immunization strategies generate durable antibody responses to SARS-CoV-2 variants that can be boosted with mRNA

Antibody responses were assessed over the period of 63 weeks. Sera was collected at week 8 (approx. peak), week 46 (memory; before mRNA boost), week 50 (peak after the mRNA boost), and week 63 (before challenge) to measure immunoglobulin G (IgG) binding to WA-1 (ancestral SARS-CoV-2 and the vaccine insert), BA.1 (ancestral Omicron) and BA.5 (**Fig. 3A-C, Fig. S5A**). At week 8, the geometric mean titer (GMT) to WA-1 in arbitrary units/mL (AU/mL) was the highest in the S-2P+S-2P group, followed by GRAd(AE)+S-2P, GRAd+S-2P and GRAd at 432740, 267183, 250180 and 79109, respectively (**Fig. 3B**). Antibody titers waned by week 46 in all groups, decreasing by 3.3-, 6.7-, 5.3- and 25.8-fold in the GRAd, GRAd+S-2P, GRAd(AE)+S-2P and S-2P+S-2P groups, respectively. The WA-1 GMT remained relatively unchanged from week 46 through week 63 in the NHP not boosted with mRNA (closed circles), while increasing significantly in NHP that did receive the mRNA boost (open circles). By week 63, the WA-1 GMT remained significantly higher in the GRAd, S-2P+S-2P and GRAd(AE)+S-2P NHP that were boosted with mRNA (GRAd: 21170 *v.* 107163; S-2P+S-2P: 17034 *v.* 155299; GRAd+S-2P: 64166 *v.* 48979; GRAd(AE)+S-2P: 50888 *v.* 131716). Antibody binding kinetics and potency to BA.5 mirrored those of WA-1, albeit on average approximately 4-fold lower (**Fig. 3C**).

**Figure 3.**
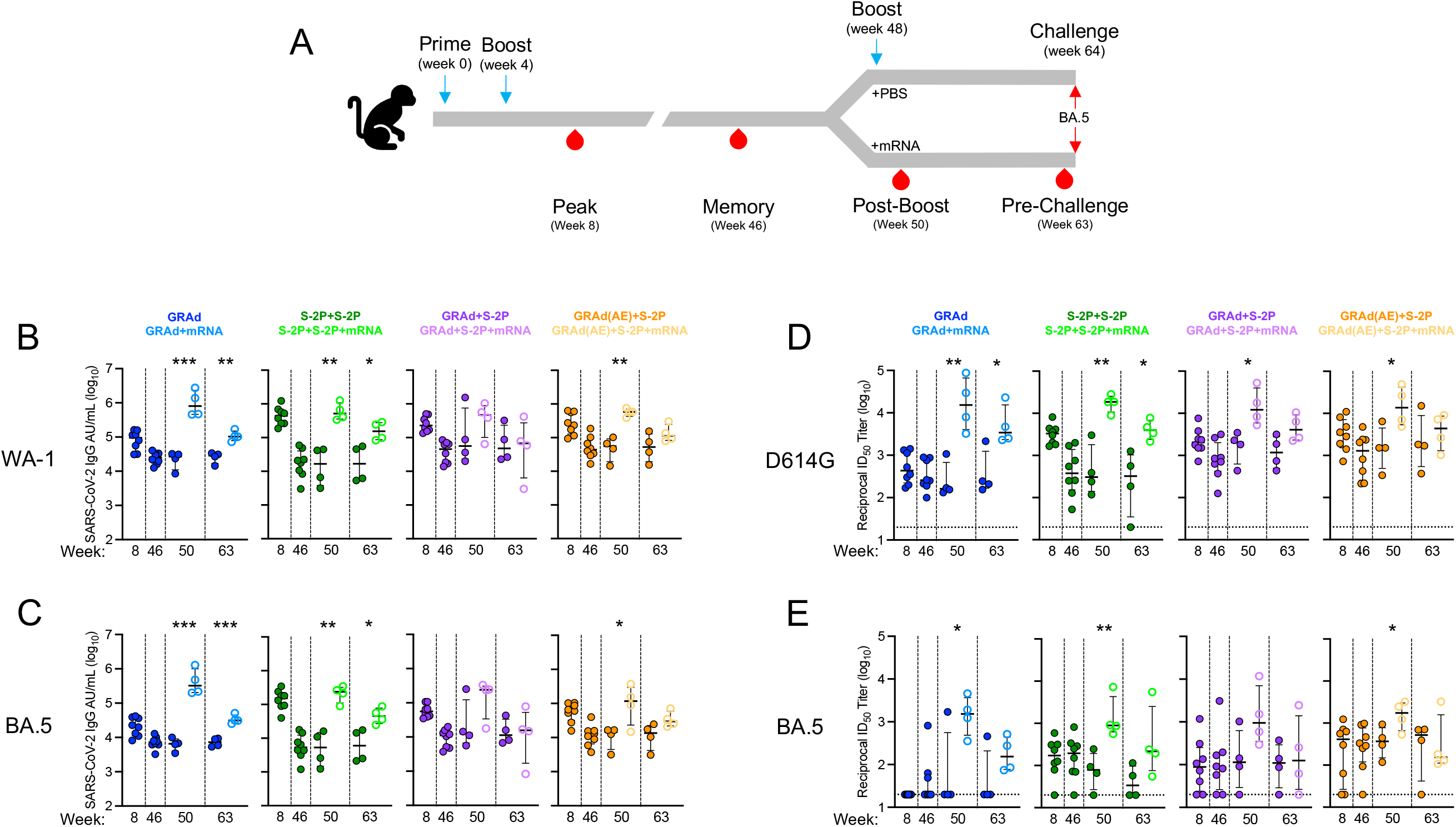
GRAd immunization strategies generate durable antibody responses to SARS-CoV-2 variants that can be boosted with mRNA. (A) Sera were collected at week 8, 46, 50 and 63. (B and C) IgG-binding titers to (B) ancestral WA-1 S and (C) BA.5 S expressed in AU/mL. (D and E) Neutralizing titers to (D) ancestral D614G lentiviral pseudovirus and (E) BA.5 lentiviral pseudovirus expressed as the reciprocal ID_50_. Circles (B–D) represent individual NHP. Error bars represent the interquartile range with the median denoted by a horizontal black line. Assay limit of detection indicated by a horizontal dotted line which may fall below the depicted range. Vertical dashed lines are for visualization purposes only. Eight immunized NHP, split into 2 cohorts of 4 NHP post mRNA boost. Statistical analysis shown for corresponding timepoints between mRNA boosted and non-boosted cohorts. * *p* <0.05, ** *p* <0.01, *** *p* <0.001. Eight immunized NHP at week 8 and 46, 4 immunized NHP at week 50 and 63. See also Figure S1 for experimental schema, Figure S5 for neutralization responses to D614G at week 2 and 4, Figure S6 for binding and neutralizing responses to BA.1 before and following challenge, and Figure S7 for binding and neutralizing responses to WA-1/D614G and BA.5 following challenge.

Neutralizing antibody (nAb) titers to D614G, BA.1, and BA.5 were quantified using a lentiviral pseudovirus neutralization assay (**Fig. 3D-E, Fig. S5C**). At week 8, nAb titers to D614G were highest in the S-2P+S-2P NHP with a GMT of 3150 reciprocal 50% inhibitory dilution (ID_50_), followed by GRAd(AE)+S-2P, GRAd+S-2P and GRAd at 3097, 2162 and 502, respectively (**Fig. 3D**). Week 8 nAb titers to BA.1 and BA.5 decreased significantly (**Fig. 3E, Fig. S5C**), dropping to 20 (limit of detection), 153, 94 and 197 in the GRAd, S-2P+S-2P, GRAd+S-2P and GRAd(AE)+S-2P against BA.5. The nAb titers at week 46 to D614G, BA.1 and BA.5 decreased from the week 8 peak but remained highest in the GRAd(AE)+S-2P group. Similar to the binding titers, nAb titers remained relatively stable from week 46 through 63 in NHP not boosted with mRNA, while the mRNA boost significantly increased them across all groups to levels above those of week 8 across all variants tested. By week 63, the D614G nAb GMT in the non-mRNA boosted NHP were 360, 226, 1172 and 2040 for GRAd, S-2P+S-2P, GRAd+S-2P, and GRAd(AE)+S-2P, respectively, significantly lower than in the mRNA boosted cohorts (4973, 4016, 4458 and 3662, respectively). The mRNA boost had the greatest impact on the GRAd and S-2P+S2P group nAb titers to BA.5 at week 63 (GRAd: 44 *v.* 173; S-2P+S-2P: 40 *v.* 345; GRAd+S-2P: 97 *v.* 166; GRAd(AE)+S-2P: 249 *v.* 301). Together, the data suggest that AE delivery of GRAd may stimulate superior durable humoral immunity to SARS-CoV-2.

We wondered whether the delivery of the first GRAd immunization had an impact on the initial humoral response as it has been suggested that viral vectors delivered mucosally could be less immunogenic compared to parenteral routes.^31^ The nAb titers to D614G at week 2 and week 4 were approximately equal after the first dose of GRAd in all the groups that received one dose of GRAd, irrespective of whether it was given IM or AE (**Fig. S6**). Only in the case of a single S-2P immunization did we observe significantly lower titers. Notably, nAb titers to BA.5 at week 2 and week 4 were below the limit of detection, highlighting the befit of a boost and the advantage of heterologous boosting.

Antibody binding and nAb titers were also quantified immediately following BA.5 challenge, at days 2, 4, 8 and 14 to assess the extent of secondary responses in immunized NHP and primary responses to BA.5 in control NHP (**Fig S5B,D, Fig. S7A-D**). In general, the primary binding and nAb response to BA.5 was more robust than that to WA-1/D614G or BA.1 and was detectable in some of the control NHP as early as day 2. All control NHP had a measurable binding and nAb titer to BA.5 by day 14. In contrast, secondary responses following challenge in the immunized animals were primarily observed in non-mRNA boosted NHP and were not evident until after day 8 following challenge.

### GRAd immunization strategies generate durable IgG and IgA responses to BA.5 in the lower and upper airway mucosa

Antibody responses to SARS-CoV-2 infection are critical for mediating protection.^28^ BAL fluid was collected at weeks 6, 46, 50 and 61, while nasal washes (NW) were collected at weeks 46, 50, and 61 to quantify antigen-specific IgG and IgA responses in the lung and nose to WA-1, BA.1 and BA.5 (**Fig. 4, Fig. S8-10**). At week 6, the BA.5 IgG GMT in the BAL was the highest in the S-2P+S-2P group at 426 AU/mL, followed by GRAd+S-2P, GRAd(AE)+S-2P and GRAd at 398, 361 and 76, respectively (**Fig. 4B**). The IgG titers waned by week 46, but remained highest in the GRAd(AE)+S-2P group. The mRNA boost significantly increased the IgG titers in all groups (week 50, closed *v.* open circles), but the responses mostly waned by week 61. However, IgG titer in the BAL remained elevated in the mRNA-boosted NHP (GRAd: 30 *v.* 105; S-2P+S-2P: 33 *v*. 227; GRAd+S-2P: 44 *v*. 130 and GRAd(AE)+S-2P: 47 *v*. 151). In contrast, antigen-specific IgA responses in BAL to BA.5 were almost exclusively detected in the GRAd(AE)+S-2P NHP, with a BA.5 IgA GMT of 150 at week 8 in this group, while below 30 in all the other groups (**Fig. 4C**). Responses waned by week 46, but the mRNA boost increased the IgA GMT in the GRAd(AE)+S-2P group from 69 to 117 at week 50, albeit non-significantly. The mRNA boost did not increase the BAL IgA GMT in any of the other groups. By week 61, antigen-specific IgA had mostly waned in all groups except in the GRAd(AE)+S-2P NHP. Together, this data suggest that AE delivery of GRAd stimulates antigen-specific IgA responses in the lung.

**Figure 4.**
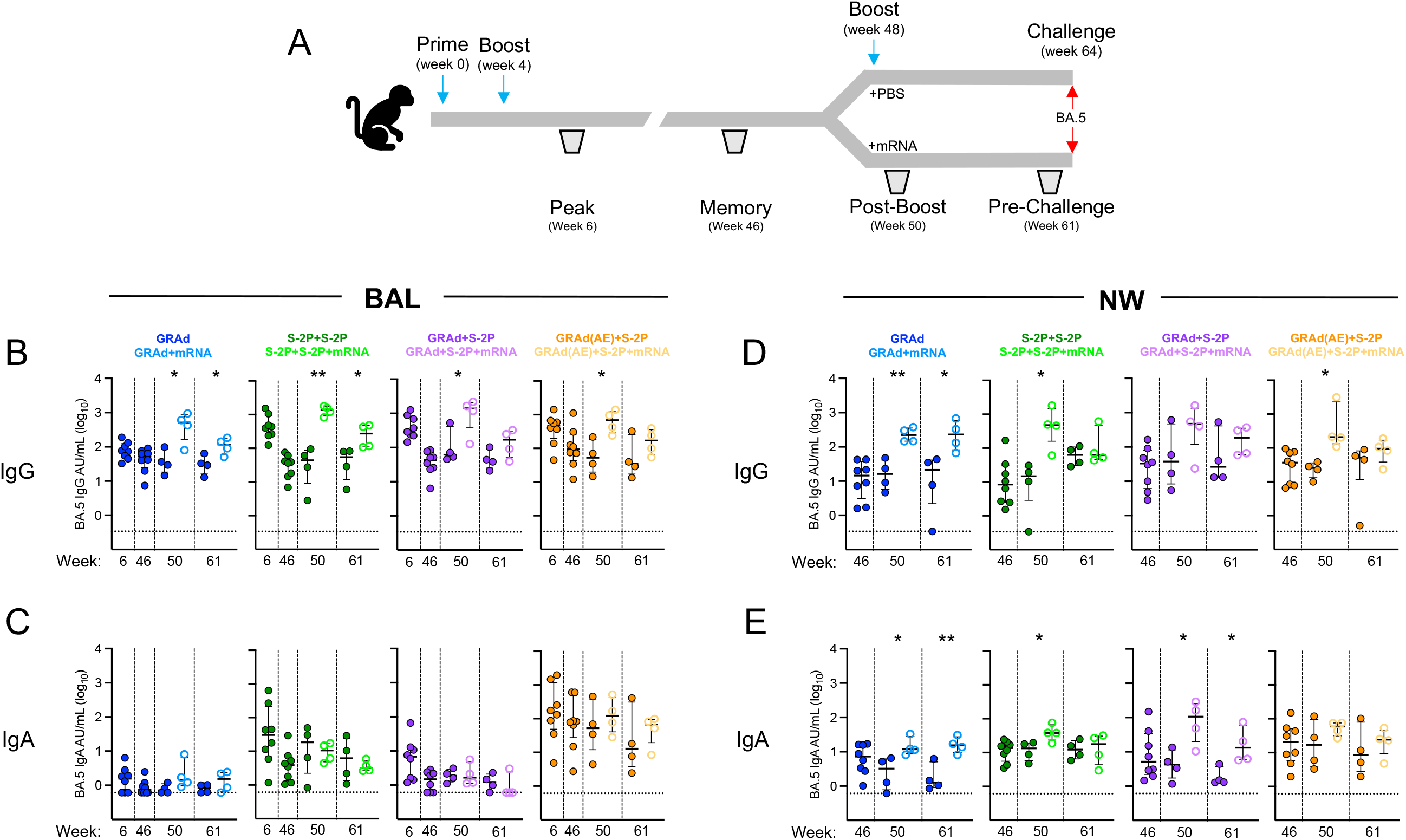
GRAd immunization strategies generate durable IgG and IgA responses to BA.5 in the lower and upper airway mucosa. (A) BAL (B and C) was collected at week 6, 46, 50 and 61 and NW (D and E) was collected at week 46, 50, and 61. (A and B) IgG (B) and IgA (C) antibody binding titers to BA.5 expressed in AU/mL in BAL. (C and D) IgG (D) and IgA (E) antibody binding titers to BA.5 expressed in AU/mL in NW. Circles (B–E) represent individual NHP. Error bars represent the interquartile range with the median denoted by a horizontal black line. Assay limit of detection indicated by a horizontal dotted line. Vertical dashed lines are for visualization purposes only. Eight vaccinated NHP, split into 2 cohorts of 4 NHP post mRNA boost. Statistical analysis shown for corresponding timepoints between mRNA boosted and non-boosted cohorts. * *p* <0.05, ** *p* <0.01. Eight immunized NHP at week 6 and 46, 4 immunized NHP at week 50 and 61. See also Figure S1 for experimental schema, Figure S8 for WA-1 and BA.1 IgG and IgA BAL and NW binding titers prior to challenge, Figure S9 for IgG and IgA binding titers in BAL following challenge and Figure S10 for IgG and IgA binding titers in NW following challenge.

In the NW, antigen-specific IgG titers to BA.5 at week 46 were approximately equal across all groups (**Fig. 4D**). The mRNA boost increased the IgG titer significantly in most groups (week 50 GMT: GRAd: 15 *v.* 221; S-2P+S-2P: 7 *v*. 399; GRAd+S-2P: 43 *v*. 302 and GRAd(AE)+S-2P: 22 *v*. 349). At week 61, the IgG titers had waned, but remained significantly elevated in the GRAd NHP that received an mRNA boost (week 61 GMT: GRAd: 8 *v*. 201; S-2P+S-2P: 58 *v*. 100; GRAd+S-2P: 44 *v*. 143 and GRAd(AE)+S-2P: 19 *v*. 79). We also detected antigen-specific IgA responses in the NW in all groups (**Fig. 4E**). At week 46, the IgA GMT was 6, 11, 8, and 18 in the GRAd, S-2P+S-2P, GRAd+S-2P and GRAd(AE)+S-2P groups, respectively. In contrast to the IgA measured in the BAL, the IgA in the NW was boosted after administration of the mRNA in most groups. Notably, however, the mRNA boost did not significantly increase the IgA titer in the NW of GRAd(AE)+S-2P NHP (week 50: GRAd: 2 *v*. 14; S-2P+S-2P: 11 *v*. 38; GRAd+S-2P: 4 *v*. 70; GRAd(AE)+S-2P: 16 *v*. 51). By week 61, NW IgA titers had waned, but generally remained elevated in NHP boosted with mRNA across the groups except in the S-2P+S-2P NHP.

We also quantified antigen-specific IgG and IgA titers against WA-1 and BA.1 in the BAL and NW (**Fig. S8**). In general, the titers and kinetics to WA-1 and BA.1 mirrored those to BA.5 but were expectedly higher. This suggests that despite having the ancestral WA-1 insert, GRAd can generate antigen-specific humoral responses in the upper and lower airway mucosa to newly emerging SARS-CoV-2 variants that have high levels of neutralization escape.^32^ Finally, we quantified IgG and IgA titers in the BAL and NW immediately following challenge (**Fig. S9**, **Fig. S10**). IgG primary responses in the control NHP were predominantly detected against BA.5, appearing after day 8 following challenge (**Fig. S9-10A-C**). In immunized NHP, evidence of a secondary antigen-specific IgG response was observed, appearing as early as 2 days following challenge in many of the groups in both the BAL and NW. As for IgA, we also observed evidence for a primary response in the control NHP and a secondary response in the immunized groups (**Fig. S9-10D-F**). Together, our mucosal antibody data suggest robust antigen-specific IgG and IgA responses are generated following immunization with GRAd.

### GRAd(AE)+S-2P generates potent year-long S-specific T cell responses in the lung

While SARS-CoV-2-specific mRNA-based vaccines elicit S-specific T_h_1, T_fh_, and CD8+ T cell responses in NHP and humans, their frequencies are relatively low.^33,34^ Furthermore, rapid control of SARS-CoV-2 may depend on fast localized T cell responses in the lung. This could be achieved by establishing a large population of SARS-CoV-2-specific T cells at the sites of infection. With some notable exceptions, all our primary immunization strategies generated T_h_1, T_fh_, and CD8+ S-specific T cell responses in the lung (**Fig. 5, Fig. S11**). The percentage of memory CD4+ T cells with a T_h_1 phenotype (IL-2, TNF and IFNγ) peaked in all groups at week 6, with GRAd, S-2P+S-2P, GRAd+S-2P, and GRAd(AE)+S-2P having 0.82, 0.65, 0.84 and 14.26, respectively, of their CD4+ T_h_1 cells positive for WA-1 S (**Fig. 5A**). Thus, on average the GRAd(AE)+S-2P NHP had approximately 17-20 times as many T_h_1 cells than the other groups at week 6. The CD4+ T_h_1 cells contracted after week 6, remaining below one percent through week 61, even after the mRNA boost, in the GRAd, S-2P+S-2P and GRAd+S-2P groups. Despite a contraction, the GRAd(AE)+S-2P NHP maintained high percentages of S-specific CD4+ T_h_1 cells, even in the absence of the mRNA boost (week 46: 7.41%; week 50: 6.87%; week 61: 6.90%). The mRNA boost did not significantly increase the percentage of S-specific CD4+ T_h_1 cells in this group (week 50: 8.35%; week 61: 4.39%). On the oppositive side of the spectrum, the percentage of S-specific memory CD4+ T_h_2 cells (IL-4 and IL-13) was below 0.2% before challenge in most NHP (**Fig. S12A**).

**Figure 5.**
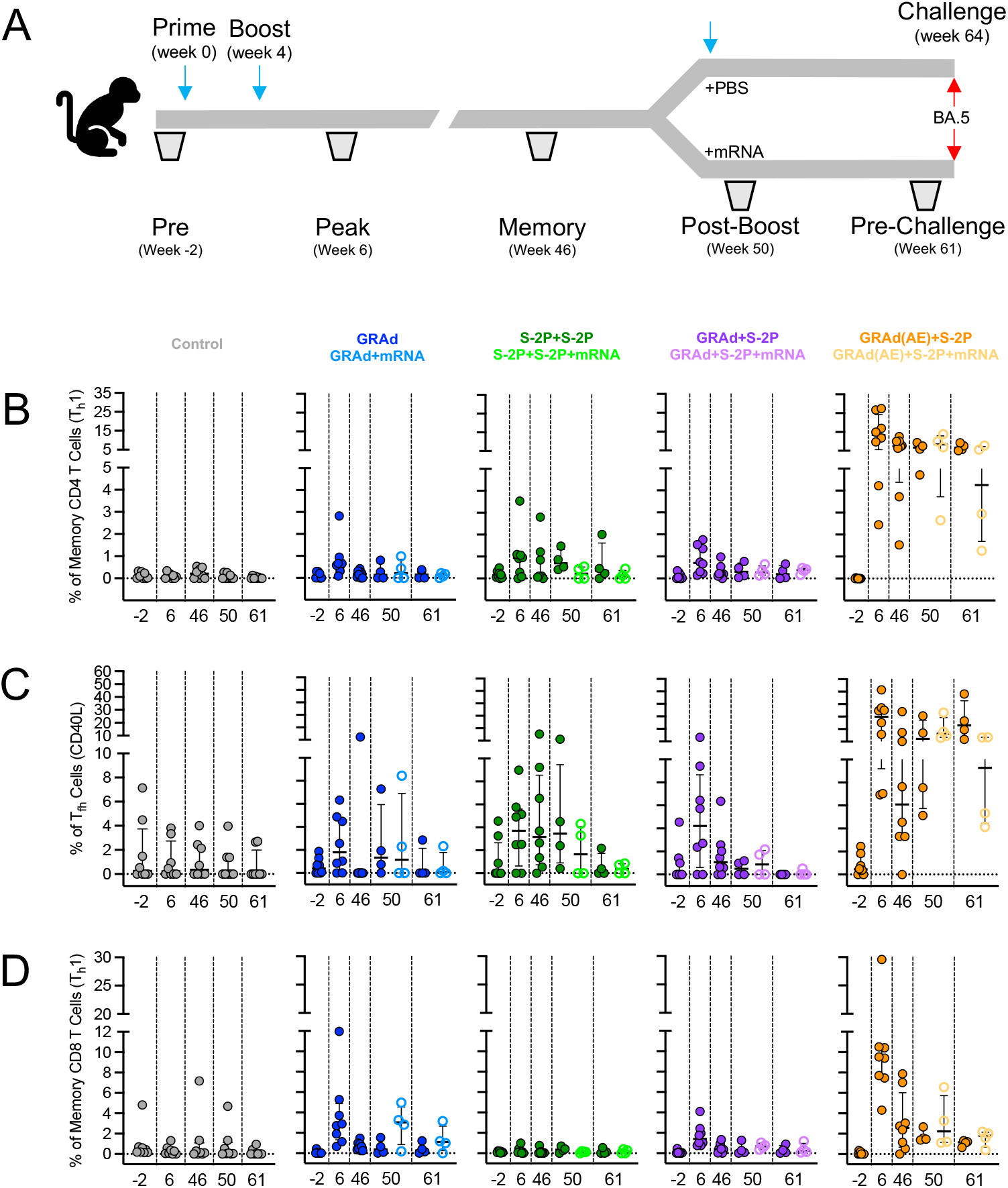
AE GRAd generates potent year-long S-specific T cell responses in the lung. (A) BAL cells were collected at week −2, 6, 46, 50 and 61. (B–D) Cells were stimulated with SARS-CoV-2 S1 and S2 peptide pools (WA-1) and then measured by intracellular cytokine staining. (A) Percentage of memory CD4+ T cells with T_h_1 markers (IL-2, TNF, or IFNγ) following stimulation. (B) Percentage of T_fh_ cells that express CD40L. (C) Percentage of CD8 T cells expressing IL-2, TNF, or IFNγ. Circles in (B–D) indicate individual NHP. Error bars represent the interquartile range with the median denoted by a horizontal black line. Dotted lines set at 0%. Reported percentages may be negative due to background subtraction and may extend below the range of the y-axis. Eight vaccinated NHP, split into 2 cohorts of 4 NHP post mRNA boost. Eight control NHP, 8 immunized NHP at week 6 and 46, 4 immunized NHP at week 50 and 61. See also Figure S1 for experimental schema, Figure S11 for T cell gating strategy, Figure S12 for T_h_2 and T_fh_ (IL-21) responses in BAL prior to and following challenges, Figure S13 for CD4+ T_h_1, T_fh_ (CD40L) and CD8+ T cell responses following challenge and Figure S14 for T cell responses in blood.

Next, we quantified the percentage of CD40L+ T_fh_ cells, which could participate in the S-specific memory B cell response.^35–37^ Much like the memory CD4+ T_h_1 cells, lung CD40L+ T_fh_ cells peaked at week 6 in all groups, though in many NHP a response was not detected (**Fig. 5B**). On average, the percent of lung CD40L+ T_fh_ cells at week 6 was 2.52%, 3.72%, 4.99% and 26.65% in the GRAd, S-2P+S-2P, GRAd+S-2P, and GRAd(AE)+S-2P groups, respectively. Lung CD40L+ T_fh_ cells contracted dramatically in the GRAd, S-2P+S-2P and GRAd+S-2P, with these groups having less than one percent CD40L+ T_fh_ positive cells at week 61, even after an mRNA boost. In contrast, at week 61, the GRAd(AE)+S-2P group had 26.51% CD40L+ T_fh_ cells in the lung. IL-21+ T_fh_ cells mirrored CD40L+ T_fh_ responses, displaying similar kinetics, but with lower overall frequency (**Fig. S12B**).

Antigen-specific memory CD8+ T cells with a T_h_1 phenotype were induced by our immunization strategies in all groups except the S-2P+S-2P group (**Fig. 5C**). This was not surprising given the inability of subunit vaccine regimens to stimulate CD8+ T cell responses.^38^ At week 6, 3.77%, 1.72% and 11.12% were S-specific memory CD8+ T cells in the GRAd, GRAd+S-2P, and GRAd(AE)+S-2P groups, respectively. The mRNA dose boosted CD8+ T cell responses in the GRAd group, but not the others, although this was not statistically significant. The CD8+ T cell responses contracted in all groups through week 61, but remained on average above 1% in the GRAd(AE)+S-2P NHP and below 1% in the other groups, regardless of whether NHP were boosted with mRNA or not.

We also quantified the S-specific T_h_1 and T_h_2, T_fh_, and CD8+ T cells following challenge (**Fig. S12C,D, Fig. S13**). Primary CD4+ T_h_1 and CD40L+ T_fh_ responses in the controls were evident at day 8, and secondary responses were also observed at day 8 in the immunized groups, particularly in the GRAd, S-2P+S-2P and GRAd+S-2P NHP that were not boosted with mRNA. Memory CD4+ T_h_1 and CD40L+ T_fh_ remained relatively unchanged from day 2 through day 14 in the GRAd(AE)+S-2P NHP. CD8+ T cells remained relatively unchanged through the challenge phase across all groups.

Finally, we asked whether the induction of potent year-long memory CD4+ T cell responses observed in the lung of GRAd(AE)+S-2P group was also present in blood. The percentage of S-specific CD4+ T_h_1 cells in blood of GRAd(AE)+S-2P NHP never rose above 1% from week 8 through week 63 (**Fig. S14A**). This observation was in sharp contrast to our observations in the lung, in which at the week 8 peak approximately 14% of the CD4+ T cells were S-specific. Despite the contraction, 3-7% of the CD4+ T cells in the lung were S-specific at week 63, in contrast to the less than 0.2% in the blood. This finding highlights the impact of AE delivery of a viral-vector on the ability to stimulate and retain antigen-specific T cells in the lung. We observed a very similar discrepancy in lung *v*. blood in the CD40L+ and IL-21+ T_fh_ cells (**Fig. S14B, E**) and S-specific memory CD8+ T cells (**Fig. S14C**), while CD4+ T_h_2 cells in the blood were undetectable in most NHP (**Fig. S14D**). Overall, our T cell data indicate potent and durable S-specific T cells are induced in the lung mucosa by GRAd(AE)+S-2P.

### Cross-reactive S-specific memory B cells are generated following immunization

Our observations of durable binding and neutralizing antibody titers in the serum and mucosa suggested an active involvement of SARS-CoV-2 S-specific cross-reactive memory B cells. To address this, we quantified the frequency of B cells in the blood specific to pairs of fluorochrome-labeled S-2P probes (**Fig. S15**), including WA-1 and BA.5 (**Fig. 6**) and WA-1 and BQ.1.1 (**Fig. S16**) before challenge at weeks 8, 46, 50, 63, and days 8 and 14 following challenge.

**Figure 6:**
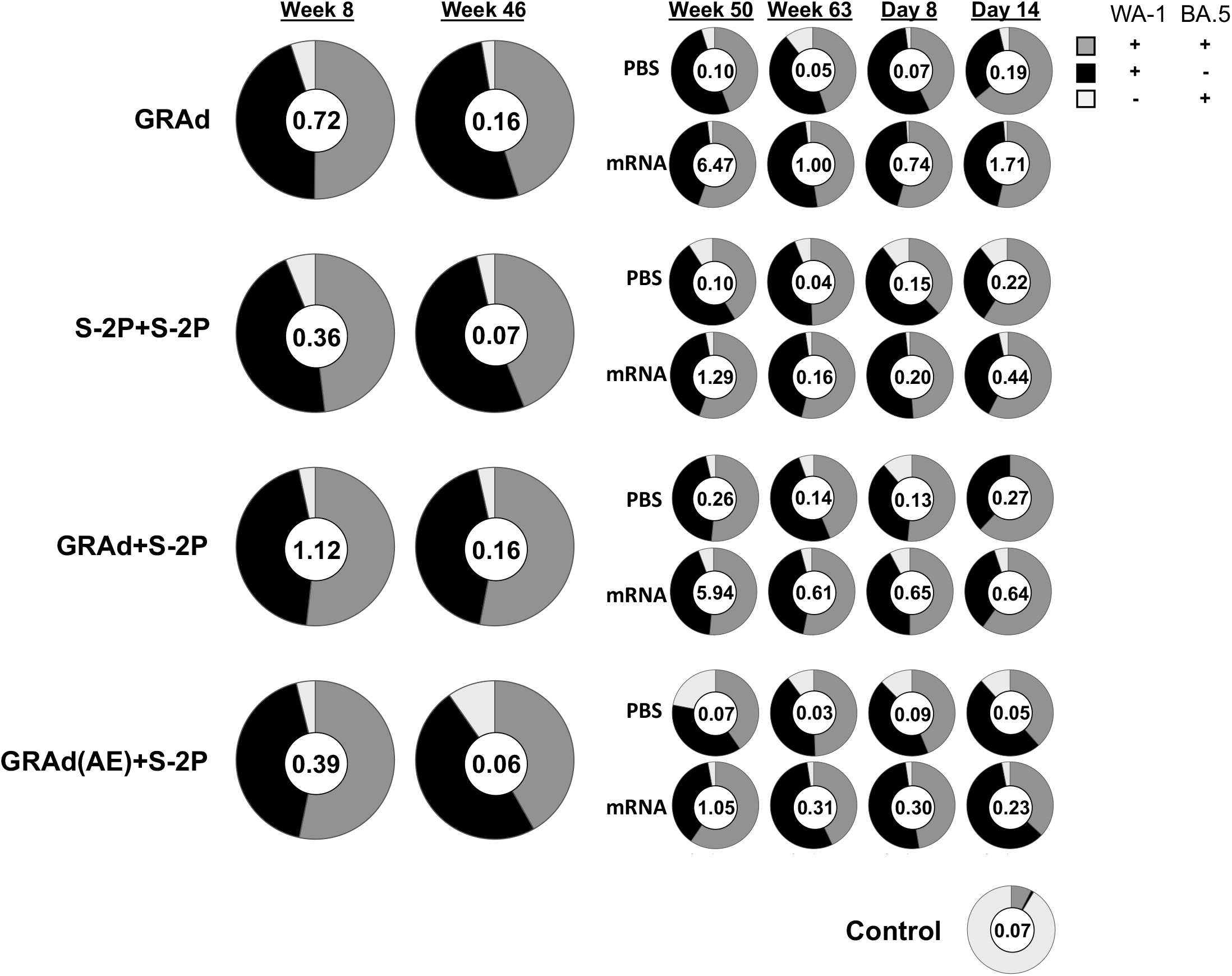
Cross-reactive S-specific memory B cells are generated following immunization. Pie charts indicate the frequency (numbered circle at the center) and proportion of total S-2P-binding memory B cells that are dual specific for WA-1 and BA.5 (dark gray), specific for WA-1 (black), or specific for BA.5 (light gray) for all NHP in each group and timepoint in the blood at week 8, 46, 50 and 63 post-immunization, and days 8 and 14 post-challenge. Seven or eight NHP per group at week 8 and 46, 3-4 NHP per group at week 50 and 63, and day 8, 1-2 NHP at day 14. See also Figure S1 for experimental schema, Figure S15 for B cell gating strategy and Figure S16 for WA-1 and BQ.1.1 cross-reactive B cell responses in blood.

At week 8 the B cell frequency (total S-2P specific memory B cells) was 0.72, 0.36, 1.12 and 0.39 percent in the GRAd, S-2P+S-2P, GRAd+S-2P, and GRAd(AE)+S-2P groups, respectively (**Fig. 6**). At week 8, the relative proportion of dual specific B cells capable of binding both WA-1 and BA.5 was approximately equal across all groups and accounted for ∼50% of the total (dark grey), while WA-1 only specific B cells (black) accounted for most of the remainder (∼45%). Only a very small percentage of the B cells were exclusively BA.5 specific (<5%; light grey). By week 46, the frequency of S-specific memory B cells had declined uniformly across the groups, decreasing on average by ∼80% from the week 8 peak, while the relative proportion of probe-specific B cells remained approximately the same.

In contrast to our T cell data, the mRNA boost had a pronounced effect on the expansion of the S-specific B cell compartment, particularly in the GRAd and GRAd+S-2P groups. At week 50, the B cell frequency was 6.47, 1.29, 5.94 and 1.05 percent in the GRAd, S-2P+S-2P, GRAd+S-2P, and GRAd(AE)+S-2P groups boosted with mRNA, respectively (**Fig. 6**), representing 3944%, 1743%, 3613% and 1650% increases from week 46. Except for the GRAd+S-2P group, the mRNA boost may have expanded the dual-specific B cell population. However, by week 63 the frequency of S-specific memory B cells had declined on average by ∼80% across all groups from week 50. Nevertheless, at week 63 the mRNA-boosted cohorts in the GRAd, S-2P+S-2P, GRAd+S-2P, and GRAd(AE)+S-2P groups had approximately 20-, 4-, 4- and 10-fold higher S-specific B cells in blood, respectively, compared to non-boosted counterparts.

Lastly, we measured the S-specific B cell response on days 8 and 14 following challenge to assess whether BA.5 challenge had expanded the B cell compartment. In control NHP, the frequency of S-specific B cells at day 14 was low, but ∼90% of the response was BA.5 specific. Evidence of a secondary response was not uniform across the groups. In the GRAd group, the B cell frequencies in the non-boosted cohort rose from 0.05 at week 63 to 0.07 on day 8 and to 0.19 on day 14, suggesting an expansion due to the challenge. A similar phenomenon was observed in the mRNA-boosted GRAd cohort, and in the S-2P+S-2P and GRAd+S-2P cohorts, regardless of whether they were boosted or not. In contrast, we did not observe an expansion in the non-boosted GRAd(AE)+S-2P cohort, going from 0.03 at week 63 to 0.09 on day 8 and 0.05 on day 14, nor in the presence of the mRNA boost, going from 0.31 at week 63 to 0.30 on day 8 to 0.23 on day 14. Additionally, this was the only group we observed a decline in the proportion of dual-specific B cells and an increase in WA-1 only specific B cells over time. Similar kinetics were observed when WA-1 and BQ.1.1 probes were used, albeit the relative proportion of dual specific B cells was lower (**Fig. S16**). Overall, all our immunizations generated durable S-specific B cell responses in blood that could be boosted, but in the absence of a boost they contracted over time.

### Priming with IM or AE GRAd alters serum antibody epitope profile in presence or absence of mRNA boost

To evaluate the impact of GRAd-or adjuvanted protein subunit-based immunogens and the impact of mRNA boosting on epitope targeting of serum antibodies to SARS-CoV-2 Spike, we used serum antibody competition assays. We determined the relative serum reactivity (as percent competition) to 18 distinct antigenic sites spanning the S1, S2, NTD and RBD subdomains of the homologous spike protein (WA-1). Serum was evaluated at week 63, immediately before BA.5 challenge.

All primary immunization strategies incorporating GRAd, both in the presence or absence of mRNA boost, showed breadth of reactivity spanning epitopes in the S2, NTD, and RBD subdomains, resembling earlier findings following mRNA-1273 vaccination (**Fig. 7A-B**).^39^ In contrast, the adjuvanted protein subunit group (S-2P+S-2P) demonstrated a distinct epitope profile characterized primarily by an absence of reactivity to 3 out of 4 antigenic sites in the NTD subdomain, as well as dominant reactivity to RBD antigenic site D (**Fig. 7A**). An average of 38.5% of all serum antibodies targeting WA-1 S were competed by the presence of Site D, defined by monoclonal Ab (mAB) A19-46.1, in this group (**Fig. 7A**).^40^ This antigenic site falls within Class II designation,^41^ indicating its ability to block the interaction of S with the human SARS-CoV-2 receptor, hACE2 (angiotensin-converting enzyme 2), and includes 2 mutations in BA.5 (N501Y, Y505H) (**Fig. 7C**). The mRNA boost did not significantly alter the epitope profile or recover reactivity to epitopes not targeted by the primary immunization regimen (**Fig. 7B**).

**Figure 7:**
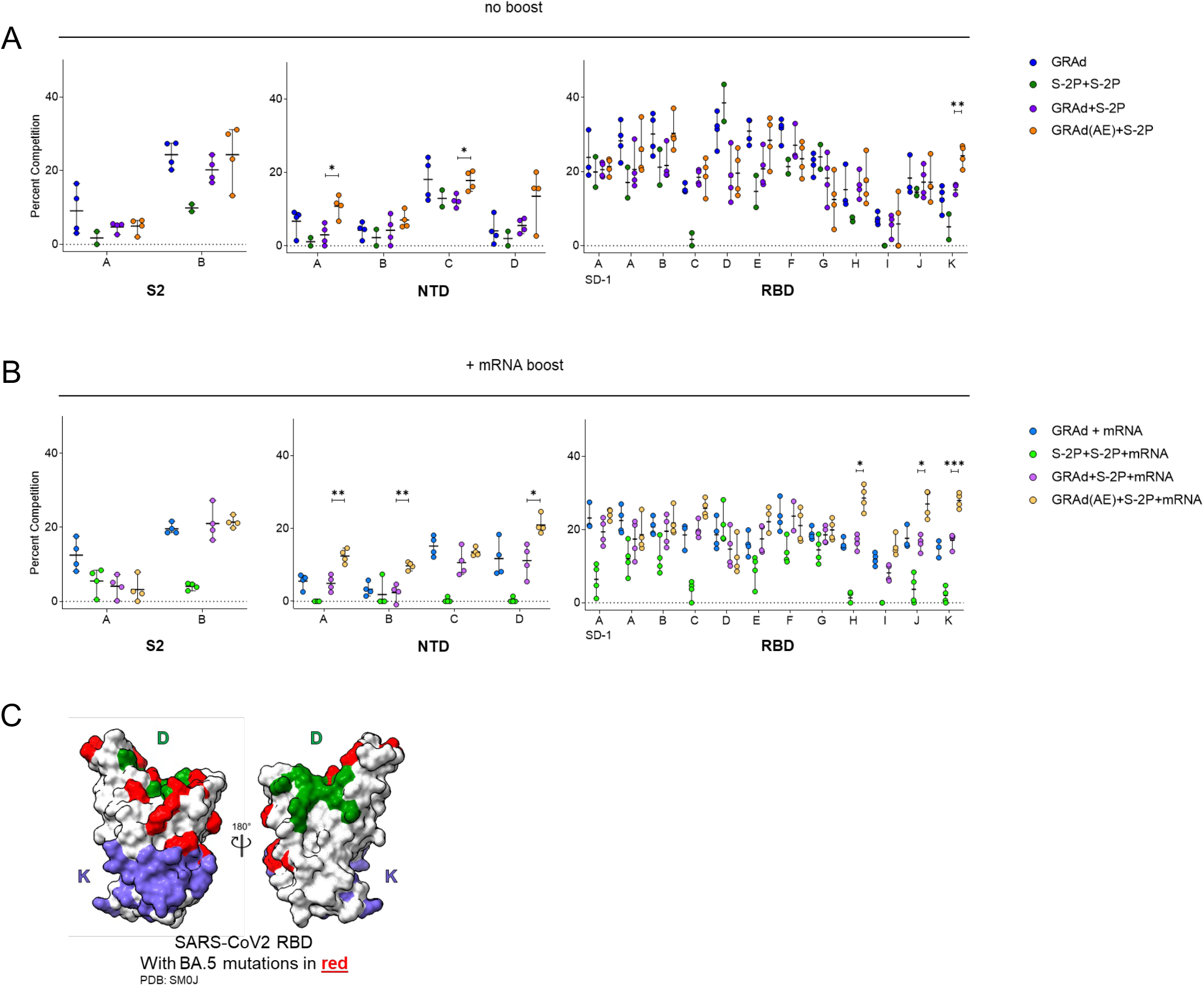
Priming with IM or AE GRAd alters serum antibody epitope profile in presence or absence of mRNA boost. (A and B) Relative serum reactivity was measured as percent of total measured serum antibody S-2P binding competed by single monoclonal antibodies (mAbs) targeting S2, NTD, and RBD epitopes on WA-1 S-2P. Relative serum reactivity was evaluated in NHPs receiving no additional boost at week 63 (panel A) or mRNA boost (panel B). Circles in (A and B) indicate individual NHP. Error bars represent the range with the median denoted by a horizontal black line. Eight vaccinated NHP, split into 2 cohorts of 4 NHP post mRNA boost. Statistical analysis shown for percentage of competition of binding to indicated epitopes at week 63 between “GRAd(AE)+S2P” and “GRAd+S2P” groups. * *p* <0.05, ** *p* <0.01, *** *p* <0.001. (C) Footprints of site D (A19-46.1) and Site K (CR3022) defining mAbs indicate areas of binding on SARS-CoV-2 RBD with BA.5 mutations highlighted in red. See also Figure S1 for experimental schema.

No significant differences in epitope targeting were observed between the GRAd and GRAd+S-2P groups, nor did these groups show significantly different epitope reactivity profiles following the mRNA boost (**Fig. 7A,B**). In contrast, NHP that received AE GRAd as the prime (GRAd(AE)+S-2P) demonstrated a remarkably different epitope profile compared to the IM counterparts (GRAd+S-2P). Immediately before challenge (week 63), GRAd(AE)+S-2P NHP had significantly higher proportions of serum reactivity to NTD antigenic site A, defining the NTD supersite, and antigenic site C compared to IM counterpart GRAd+S-2P (**Fig. 7A**). In addition, we observed a significantly higher proportion of serum antibodies targeting RBD antigenic site K (**Fig. 7A**). This site is defined by the Class IV mAb CR3022, which is highly conserved across SARS-CoV and SARS-CoV-2.^41,42^ In BA.5, this footprint contains two mutations located on the periphery of the binding site (S371F, S373P) (**Fig. 7C**). These significant differences in epitope targeting between AE and IM delivery groups were also evident after the mRNA boost (**Fig. 7B**). Together, these data highlight the impact of antigen delivery, the importance of the initial vaccination formulation and route, and the impact of the immunization series on epitope targeting and emphasizes that the initial vaccination is critical in shaping the serum antibody epitope profile following subsequent vaccinations.

## Discussion

SARS-CoV-2 vaccines that prevent or mitigate infection and transmission are urgently needed. While the currently approved vaccines have successfully curbed morbidity and mortality from SARS-CoV-2, thousands continue to die due to COVID-19 worldwide every week.^1^ An obvious tactic to reduce the number of cases, and consequently the number of deaths, would be to reduce person-to-person transmission. One approach may be with vaccines that directly stimulate local respiratory tract immunity in addition to systemic immunity. While pre-clinical research studies with intranasal (IN) vaccines have demonstrated efficacy against SARS-CoV-2, it remains unclear if this delivery method would help curtail transmission.^14–16^ It has been reported that AE delivery generates stronger immune responses than IN delivery, leading us to speculate that AE may outperform IN delivery at curtailing transmission.^43,44^ In this study, we evaluated the GRAd viral vector vaccine platform expressing pre-fusion stabilized S as a single dose, and as a heterologous prime and boost regimen that included intramuscular and AE delivery in NHP. A single GRAd dose provided over one-year protective immunity against SARS-CoV-2. Boosting with adjuvanted S-2P (GRAd+S-2P) accelerated viral clearance in the lower and upper airways. Although aerosol-delivered GRAd (GRAd(AE)+S-2P) only modestly improved protection, mRNA boosting proved more effective in this group, achieving total virus clearance in the upper airway by day 4 post-infection. GrAd vaccination triggered strong systemic and mucosal antibody responses to diverse SARS-CoV-2 variants, while GRAd(AE)+S-2P uniquely generated enduring T cell responses in the lung.

In a first-in-human, dose-escalation phase 1 trial, a single intramuscular dose of the GRAd vector expressing S-2P (abbreviated GRAd-COV2 when used in humans) was shown to be safe and immunogenic in younger (18 to 55 years old) and older (65 to 85 years old) adults.^45^ The observed favorable tolerability was confirmed and extended in a randomized, double-blinded, placebo-controlled phase 2 study, where a two GRAd-COV2 dose regimen with an interval of 21 days was also evaluated.^46^ The safety profile was similar to other viral vector vaccines,^22,47–50^ and adverse events were milder than those reported for nanoparticle encapsulated mRNA and adjuvanted subunit/protein vaccines.^34,51,52^ In both phase 1 and 2 studies, the vaccine was immunogenic after a single dose, inducing rapid binding and neutralizing antibodies that were boosted by the administration of a second GRAd-COV2 dose. The vaccine also induced potent, broad, durable cross-reactive T_h_1 S-specific CD4+ and CD8+ T cells with high proliferative capacity, that was not expanded by a homologous boost.^46,53^ When compared in SARS-CoV-2 International Units (IU), GRAd-COV2 induced binding and neutralizing antibody levels that were equivalent to those of other single-dose viral vector-based vaccines such as Ad26 and ChAdOx1,^22,34,54–56^ but lower than those reported for the two-dose regimen of mRNA vaccines.^16,57,58^ Despite the human immunogenicity data, GRAd-COV2 has not been formally evaluated for efficacy and it remains unknown how this platform would perform in a heterologous prime and boost immunization strategy, but observations from phase 1 and 2 clinical studies revealed compelling immune responses when GRAd-COV2 recipients were boosted with mRNA;^46,59^ or if the initial GRAd-COV2 dose was delivered directly into the upper and lower airway mucosa to stimulate systemic and local mucosal immunity in the respiratory tract.

In the present study in NHP, a single IM dose of GRAd given 64 weeks before challenge protected against SARS-CoV-2 BA.5 in the lung, while two IM doses of adjuvanted S-2P could not. A heterologous IM boost to IM GRAd with adjuvanted S-2P at week 4 conferred additional protection in the lung, but this was not statistically significant, while heterologous AE GRAd prime with an IM S-2P boost did not yield a meaningful protective benefit to the IM route in the lung. The mRNA boost 16 weeks before challenge enhanced protection in the lung, but this was limited to the single dose of GRAd and the S-2P+S-2P groups. In contrast, the AE prime with GRAd proved most beneficial in terms of protective efficacy in the nasal passage. Neither the single IM GRAd nor the IM S-2P+S-2P were effective at curtailing BA.5 in the nose, while the IM GRAd+S-2P and GRAd(AE)+S-2P began clearing the virus by day 4 and 2, respectively, demonstrating rapid clearance in the nose can be achieved even 60 weeks after immunization. The mRNA boost had its most pronounced effect on the GRAd(AE)+S-2P group, accelerating clearance of the virus and clearing it completely by day 4. Altogether, the protection data demonstrate that GRAd is a suitable vaccine platform against severe disease from SARS-CoV-2, and when combined with AE delivery may help curtail its transmission.

Serologically, all our vaccine regimens efficiently induced systemic binding antibody responses to the vaccine-matched SARS-CoV-2 WA-1 strain, as well as BA.1 and BA.5 variants that lasted for at least 63 weeks, even in the absence of the mRNA boost. Serum antibodies in all groups also efficiently neutralized the WA-1 strain, however, neutralizing activity against BA.1 and BA.5 weakened significantly in the GRAd and S-2P+S-2P groups over time, while the median BA.5 titer in the GRAd+S-2P and GRAd(AE)+S-2P remained stable over the 63 weeks. The mRNA boost rescued low BA.5 neutralizing antibody titers in some groups, but the titers began to wane shortly thereafter, suggesting a limited benefit of continuous boosting if the primary immunization or immunization series is inadequate. In addition, all immunization strategies efficiently induced durable mucosal IgG in the upper and lower airway to WA-1, BA.1 and BA.5. In contrast, IgA was more prevalent in the BAL of GRAd(AE)+S-2P NHP and remained elevated over 61 weeks. Surprisingly, nasal IgA was only slightly higher in the GRAd(AE)+S-2P NHP compared to the other groups, suggesting that IgA may not be playing a major role in protection against SARS-CoV-2. Indeed, while selective IgA deficiency is a common primary immunodeficiency disorder,^60^ it remains unclear whether this is a risk factor of SARS-CoV-2.^61–63^ Thus, despite all our immunization strategies inducing systemic and mucosal binding and neutralizing antibody responses, the heterologous primary vaccination strategies were superior, and AE delivery of GRAd conferred the additional benefit of mucosal humoral immune stimulation.

All our immunization strategies induced S-specific memory CD4+ T cells, and except for in the S-2P+S-2P group, memory CD8+ T cells in the blood and the lung. Like immunization with other SARS-CoV-2 vaccines, the memory S-specific T cell response in our immunization regimens skewed toward the T_h_1 phenotype, expressing IFNγ, TNF and IL-2.^27,64,65^ In the lung, the T_h_1 CD4+ T cells peaked at approximately week 6 and mostly contracted to basal levels by week 61 in all groups except in the NHP that received GRAd(AE)+S-2P, suggesting that priming the mucosa with GRAd resulted in the generation of robust and long-lived T cells in the lung, an observation that was not reproduced in the blood, highlighting the importance of directly stimulating the lung mucosa.^66^ Indeed, IM mRNA and viral-vectored, including GRAd, SARS-CoV-2 vaccines induce weak, if any, S-specific CD4+ and CD8+ T cell responses in the airways of NHP.^67^ In our study, the mRNA dose did not boost the CD4+ T cell responses in any group, again suggesting there may be a limited benefit of periodic IM boosters if the primary immunization did not stimulate them in the first place. Memory S-specific CD8+ T cells that expressed IFNγ, TNF and IL-2 were also more prevalent in the lung of GRAd(AE)+S-2P NHP, but this population contracted through week 61, in contrast to the CD4+ T cell population.

Finally, we found similar kinetics in the T_fh_ populations in the GRAd(AE)+S-2P group, an important observation as previous studies suggest they contribute to long-term immunity and are associated with protection against respiratory disease.^68–71^ Together, the data suggest that AE delivery of GRAd can stimulate robust, antigen-specific long-lived memory CD4+ T cells in the mucosa, a clear advantage as mucosal immune responses at the site of infection are essential.^72^

Antigen-specific B cells that bound S of multiple SARS-CoV-2 variants, including WA-1, BA.1 (data not shown), BA.5, and BQ.1.1 were also observed in all our immunization strategies. While most of the B cells were dual specific for WA-1 and BA.5, we observed a notable loss of dual specific WA-1 and BQ.1.1 B cells, and this loss was accompanied by a relative increase in WA-1 only specific B cells, suggesting that the WA-1 imprint on immune memory may continue to weaken against new SARS-CoV-2 variants and may be difficult to overcome.^3^ However, it remains unknown how a matched variant boost would affect the B cell specificities in this study. Nevertheless, the relative proportion of cross-reactive B cells remained stable for over a year, and while the mRNA boost was able to increase the relative proportion of these cross-reactive B cells in some groups, the effect appears to have been only transient. Although the mRNA boost did expand the B cell frequency to a greater extent in the GRAd and GRAd+S-2P groups compared to the S-2P+S-2P and GRAd(AE)+S-2P groups, a contraction quickly followed, a phenomenon that appears to be more substantial in NHP than in humans.^73^ The differences we observed in B cell frequencies between the GRAd+S-2P and GRAd(AE)+S-2P, a change in only GRAd priming via IM or AE, led us to explore whether this alteration affected epitope specificities to S. The fact that we observed differences in epitope targeting by only changing the delivery of GRAd from IM to AE, and that epitopes targeted by AE delivery appear to be more conserved, suggests that the mechanism of antigen presentation that occurs when antigens are delivered to the lung differs from when delivered to the lymph nodes via the musculature. This is entirely plausible given the anatomical differences between muscle and lung immune cells, and the lymph nodes each tissue drains to.^74,75^ Thus, delivery of the immunizing agent to the anatomical site in which the pathogen causes disease may provide greater immune fidelity.

In summary, GRAd based vaccines encoding S-2P were highly immunogenic and protective against SARS-CoV-2 BA.5 in macaques and would have likely been effective against earlier variants. The efficacy and immunogenicity of GRAd was improved via heterologous boost with S-2P, while protection with S-2P+S-2P was inadequate. Boosting with mRNA yielded transient increases in systemic and mucosal antibody binding and neutralizing responses and overall B cell frequencies, but it did not have a meaningful impact on T cell responses. Our data support the further development of the GRAd platform for use as a prime only or as part of a prime and boost combination with other approved SARS-CoV-2 vaccines.

### Limitations of the study

In our study, the immunogens (GRAd viral vector and S-2P protein) were derived from the ancestral Wuhan-Hu-1/USA-WA1/2020, while the NHP were challenged with the SARS-CoV-2 BA.5, a significant shift from vaccine insert to challenge virus. While we detected binding and neutralizing antibody responses to BA.5, it is possible that GRAd efficacy could be improved if the insert was matched to the challenge virus. In that vein, T cell responses reported in this study were measured against WA-1, and it is possible that antigenic differences between WA-1 and BA.5 could have resulted in weaker BA.5-specific T cell responses. Thus, Omicron sublineage-specific T cell responses should be evaluated in future studies. Furthermore, we did not evaluate the immunogenicity and efficacy of a single dose of AE GRAd, and while the S-2P boost likely contributed to the observed protection in the GRAd(AE)+S-2P group, we would likely have observed similar humoral and cellular responses and protection in the absence of a boost in this group. Finally, while we evaluated the efficacy against BA.5, it was subsequently displaced by BQ.1.1 and it by XBB.1.5. It is unknown how our immunogens and vaccination strategies would fare upon challenge from currently circulating Omicron sub-lineage variants, but like bivalent mRNA vaccines, viral-vectors could be formulated as a mixture of viruses that encode S from various SARS-CoV-2 variants.

## Methods

### Resource availability

#### Lead contact

Further information and requests for resources should be directed to and will be fulfilled by the lead contacts, Robert A. Seder Nancy J. Sullivan.

#### Materials availability

This study did not generate new unique reagents.

#### Data and code availability

- All data reported in this paper will be shared by the lead contact upon request.
- This paper does not report original code.
- Any additional information required to reanalyze the data reported in this paper is available from the lead contact upon request.

### Experimental model and subject details

#### Rhesus macaque model and immunizations

All experiments were conducted according to NIH regulations and standards on the humane care and use of laboratory animals as well as the Animal Care and Use Committees of the NIH Vaccine Research Center and BIOQUAL, Inc. (Rockville, Maryland). All studies were conducted at BIOQUAL, Inc. Forty, four-to nine-year-old rhesus macaques of Indian origin were stratified into five groups of eight based on sex, age, and weight. Group one was immunized with 5×10^10^ GRAd32-Gag at week 0 and placebo (phosphate-buffered saline - PBS) at week 4. This group served as the control. Group two was immunized with 5×10^10^ GRAd32-S-2P at week 0 and placebo at week 4. Group three was immunized 5 μg adjuvanted S-2P (750 μg alum (aluminum hydroxide) and 1500 μg CpG 1018 (Dynavax Technologies); the same adjuvants were used when applicable) at week 0 and week 4. Group four was immunized with 5×10^10^ GRAd32-S-2P at week 0 and with 5 μg adjuvanted S-2P at week 4. Group five was immunized with 5×10^10^ GRAd32-S-2P delivered via aerosol (AE) at week 0 and with 5 μg adjuvanted S-2P at week 4. AE was delivered in a 2 ml volume via a pediatric silicon face mask (PARI SMARTMASK^®^ Baby/Kids) attached to an Investigational eFlow Nebulizer System (PARI Respiratory Equipment, Inc., Midlothian, VA, USA) that delivered 4 μM particles into the lung, as previously described.^76^ At week 48 (∼ 44 weeks after the second immunization), the eight macaques in each group were subdivided into two groups of 4 (except group one) and boosted with 30 μg BNT162b2. The week 0 and week 4 immunizations were delivered intramuscularly in 1 mL diluted in PBS (except AE GRAd32-S-2P) into the right deltoid, while the week 48 immunizations were delivered intramuscularly in 1 mL diluted in PBS into the right quadricep.

### Method details

#### Cells and viruses

VeroE6-TMPRSS2 cells were generated at the Vaccine Research Center, NIH, Bethesda, MD. Viruses were propagated in Vero-TMPRSS2 cells to generate viral stocks. Viral titers were determined by focus-forming assay on VeroE6-TMPRSS2 cells. Viral stocks were stored at - 80°C until use.

#### GRAd32 vectors expressing SARS-CoV-2 S-2P

The GRAd32 vector was isolated, amplified, classified and constructed as a vector as previously described in detail.^17^ The GRAd32 vector expressing SARS-CoV-2 pre-fusion stabilized Spike (S-2P) was generated as previously described.^17^ Briefly, the SARS-CoV-2 S-2P gene was generated by subcloning a human codon-optimized version of the SARS-CoV-2 S into a shuttle plasmid between the AscI and PacI restriction sites. Two mutations were introduced to convert the 986 lysine and 987 valine (KV) amino acids (aa) into prolines (PP or 2P) to stabilize the protein in its pre-fusion state.^77^ A hemagglutinin (HA) tag was fused downstream of the last SARS-CoV-2 S protein aa (Threonine1273) flanked at its 5′ and 3′ side by a Glycine and a Serine, respectively, to facilitate antigen expression detection. A minimal Kozak sequence (5′-CCACC-3′) was placed immediately upstream of the start codon to enable efficient initiation of translation. The cassette encoding for the SARS-CoV-2 S-2P was inserted by homologous recombination in the E1 locus of the GRAd32 vector. All cloning PCR amplifications were performed using the Q5 High-Fidelity DNA Polymerase (New England Biolabs) according to standard procedures. The GRAd32 S-2P vector was expanded in a 2 L Bioreactor (Biostat B DCU; Sartorius). The titer of virus contained in the bulk cell lysates was quantified by qPCR. Extensive purification of GRAd-S-2P vector was obtained by applying an orthogonal chromatographic method.

#### Expression and purification of SARS-CoV-2 S-2P

SARS-CoV-2 S-2P protein (S2P7471) was produced as previously described.^78^ Briefly, S2P7471 was expressed and purified from the CHO-DG44 cell line. S2P7471 was clarified through centrifugation and Satopore XLG 0.8/0.2 filters. S2P7471 was concentrated and buffer exchanged by tangential flow filtration into 1x PBS, purified via Nickle NTA nitrilotriacetic acid chromatography, and purified by Superose 6 size-exclusion chromatography (SEC). Finally, S2P7471 was cleaved by HRV3C protease, and the cleaved product was loaded onto a Superose 6 SEC column. Peak fractions of S2P7471 from the SEC were pooled, concentrated using SPIN-X UF spin filters, spiked to 5% sucrose and sterile filtered. S2P7471 was concentrated to 0.607 mg/mL, flash-frozen in liquid nitrogen and maintained at −80 °C until use.

#### Sequencing of BA.5 virus stock

NEBNext Ultra II RNA Prep reagents and multiplex oligos (New England Biolabs) were used to prepare Illumina-ready libraries, which were sequenced on a NextSeq 2000 sequencer (Illumina). Demultiplexed sequence reads were analyzed in the CLC Genomics Workbench v.22.0.2 by (1) trimming for quality, length, and adaptor sequence, (2) mapping to the Wuhan-Hu-1 SARS-CoV-2 reference (GenBank no. NC_045512), (3) improving the mapping by local realignment in areas containing insertions and deletions (indels), and (4) generating both a sample consensus sequence and a list of variants. Default settings were used for all tools.

#### Viral challenge

Macaques were challenged at week 64 with a total dose of 8 × 10^5^ PFUs of SARS-CoV-2 BA.5 kindly provided by M. Suthar (Emory). The viral inoculum was administered as 6 ×10^5^ PFUs in 3 mL intratracheally and 2 ×10^5^ PFUs in 1 mL intranasally in a volume of 0.5 mL into each nostril.

#### Serum and mucosal antibody titers

Quantification of antibodies in the blood and mucosa was performed using multiplex electrochemiluminescence serology assays by Meso Scale Discovery Inc. (MSD) as previously described.^79^ Briefly, total IgG and IgA antigen-specific antibodies to variant SARS-CoV-2 S-were determined by MSD V-Plex SARS-CoV-2 Panel 24 (K15575U, K15577U), Panel 27 (K15606U, K15608U) and Spike Panel 1 Kit (K15651U, K15653U) for S according to manufacturer’s instructions, except 25μl of sample and detection antibody were used per well. Heat inactivated plasma was initially diluted 1:1000 and 1:5000 using Diluent 100. BAL fluid and nasal washes were concentrated 10-fold with Amicon Ultra centrifugal filter devices (Millipore Sigma). Concentrated samples were diluted 1:10 and 1:100 using Diluent 100. Arbitrary units per milliliter (AU/mL) were calculated for each sample using the MSD reference standard curve. For each sample, the data reported was for diluted samples that returned results between the upper and lower limits of quantification.

#### Lentiviral pseudovirus neutralization

Neutralizing antibodies in serum or plasma were measured in a validated pseudovirus-based assay as a function of reductions in luciferase reporter gene expression after a single round of infection with SARS-CoV-2 spike-pseudotyped viruses in 293T/ACE2 cells (293T cell line stably overexpressing the human ACE2 cell surface receptor protein, obtained from Drs. Mike Farzan and Huihui Mu at Scripps) as previously described.^57,80^ SARS-CoV-2 Spike-pseudotyped virus was prepared by transfection in 293T/17 cells (human embryonic kidney cells in origin; obtained from American Type Culture Collection, cat. No. CRL-11268) using a lentivirus backbone vector, a spike-expression plasmid encoding S protein from Wuhan-Hu-1 strain (GenBank no. MN908947.3) with a p.Asp614Gly mutation, a TMPRSS2 expression plasmid, and a firefly Luc reporter plasmid. For pseudovirus encoding the S from B.1.1.529 (BA.1) and BA.5, the plasmid was altered via site-directed mutagenesis to match the S sequence to the corresponding variant sequence as previously described.^39,67^ A pre-titrated dose of pseudovirus was incubated with eight serial 5-fold dilutions of serum samples (1:20 start dilution) in duplicate in 96-well 384-well flat-bottom tissue culture plates (Thermo Fisher, cat. no. 12-565-344) for 1 hr at 37°C before adding 293T/ACE2 cells. One set of 14 wells received cells + virus (virus control) and another set of 14 wells received cells only (background control), corresponding to technical replicates. Luminescence was measured after 66-72 hr of incubation using Britelite-Plus luciferase reagent (Perkin Elmer, cat. no. 6066769). Neutralization titers are the inhibitory dilution of serum samples at which relative luminescence units (RLUs) were reduced by 50% (ID_50_) compared to virus control wells after subtracting background RLUs. Serum samples were heat-inactivated for 30-45 min at 56°C before assay.

#### B cell probe binding

Kinetics of S-specific memory B cells responses were determined as previously described.^33^ Briefly, cryopreserved PBMC were thawed and stained with the following antibodies (monoclonal unless indicated): IgD FITC (goat polyclonal, Southern Biotech), IgM PerCP-Cy5.5 (clone G20-127, BD Biosciences), IgA Dylight 405 (goat polyclonal, Jackson Immunoresearch Inc), CD20 BV570 (clone 2H7, Biolegend), CD27 BV650 (clone O323, Biolegend), CD14 BV785 (clone M5E2, Biolegend), CD16 BUV496 (clone 3G8, BD Biosciences), CD4 BUV737 (clone SK3, BD Biosciences), CD19 APC (clone J3-119, Beckman), IgG Alexa 700 (clone G18-145, BD Biosciences), CD3 APC-Cy7 (clone SP34-2, BD Biosciences), CD38 PE (clone OKT10, Caprico Biotechnologies), CD21 PE-Cy5 (clone B-ly4, BD Biosciences), and CXCR5 PE-Cy7 (clone MU5UBEE, Thermo Fisher Scientific). Stained cells were then incubated with streptavidin-BV605 (BD Biosciences) labeled BA.1 S-2P, streptavidin-BUV661 (BD Biosciences) labeled WA-1 S-2P and streptavidin-BUV395 (BD Biosciences) labeled BA.5 or BQ.1.1 S-2P for 30 minutes at 4°C (protected from light). Cells were washed and fixed in 0.5% formaldehyde (Tousimis Research Corp) before data acquisition. Aqua live/dead fixable dead cell stain kit (Thermo Fisher Scientific) was used to exclude dead cells. All antibodies were previously titrated to determine the optimal concentration. Samples were acquired on a BD FACSymphony cytometer and analyzed using FlowJo version 10.7.2 (BD, Ashland, OR).

#### Intracellular cytokine staining

Intracellular cytokine staining was performed as previously described.^33,81,82^ Briefly, cryopreserved PBMC and BAL cells were thawed and rested overnight in a 37°C/5% CO_2_ incubator. The following morning, cells were stimulated with SARS-CoV-2 S protein (S1 and S2, matched to vaccine insert) peptide pools (JPT Peptides) at a final concentration of 2 μg/ml in the presence of 3 mM monensin for 6 h. The S1 and S2 peptide pools were comprised of 158 and 157 individual peptides, respectively, as 15 mers overlapping by 11 amino acids in 100% DMSO. Negative controls received an equal concentration of DMSO instead of peptides (final concentration of 0.5 %). The following monoclonal antibodies were used: CD3 APC-Cy7 (clone SP34-2, BD Biosciences), CD4 PE-Cy5.5 (clone S3.5, Invitrogen), CD8 BV570 (clone RPA-T8, BioLegend), CD45RA PE-Cy5 (clone 5H9, BD Biosciences), CCR7 BV650 (clone G043H7, BioLegend), CXCR5 PE (clone MU5UBEE, Thermo Fisher), PD-1 BUV737 (clone EH12.1, BD Biosciences), ICOS PE-Cy7 (clone C398.4A, BioLegend), CD69 ECD (cloneTP1.55.3, Beckman Coulter), IFNγ Ax700 (clone B27, BioLegend), IL-2 BV750 (clone MQ1-17H12, BD Biosciences), IL-4 BB700 (clone MP4-25D2, BD Biosciences), TNF-FITC (clone Mab11, BD Biosciences), IL-13 BV421 (clone JES10-5A2, BD Biosciences), IL-17 BV605 (clone BL168, BioLegend), IL-21 Ax647 (clone 3A3-N2.1, BD Biosciences), and CD154 BV785 (clone 24-31, BioLegend). Aqua live/dead fixable dead cell stain kit (Thermo Fisher Scientific) was used to exclude dead cells. All antibodies were previously titrated to determine the optimal concentration. Samples were acquired on a BD FACSymphony flow cytometer and analyzed using FlowJo version 10.8.0 (BD, Ashland, OR).

#### Subgenomic RNA quantification

sgRNA was isolated and quantified by researchers blinded to vaccine status as described, ^27^ with the sole exception of the use of a new probe listed below. Briefly, total RNA was extracted from BAL fluid and nasal swabs using RNAzol BD column kit (Molecular Research Center). PCR reactions were conducted with TaqMan Fast Virus 1-Step Master Mix (Applied Biosystems), forward primer in the 5’ leader region and gene-specific probes and reverse primers as follows: sgLeadSARSCoV2_F: 5’-CGATCTCTTGTAGATCTGTTCTC-3’; N gene: N2_P: 5’-FAM-CGATCAAAACAACGTCGGCCCC-BHQ1-3’, wtN_R: 5’-GGTGAACCAAGACGCAGTAT-3’. Amplifications were performed with a QuantStudio 6 Pro Real-Time PCR System (Applied Biosystems). The assay lower LOD was 50 copies per reaction.

#### Median Tissue Culture Infectious Dose (TCID_50_) assay

TCID_50_ was quantified as previously described.^83^ Briefly, 25,000 Vero-TMPRSS2 cells were plated per well in DMEM+10% FBS + gentamycin and incubated at 37 °C, 5% CO_2_ overnight. The following day, BAL samples were serially diluted, and the plates were incubated at 37 °C, 5.0% CO_2_ for four days. Positive (virus stock of known infectious titer in the assay) and negative (medium only) control wells were included in each assay setup. The cell monolayers were visually inspected for cytopathic effect. TCID_50_ values were calculated using the Reed–Muench formula.

#### Histopathology and immunohistochemistry

Routine histopathology and detection of SARS-CoV-2 virus antigen via immunohistochemistry was performed as previously described.^83^ Briefly, 8 to 9 days and 14 to 15 days following BA.5 challenge animals were euthanized, and lung tissue was processed and stained with hematoxylin and eosin for pathological analysis or with a rabbit polyclonal anti-SARS-CoV-2 antibody (GeneTex, GTX135357) at a dilution of 1:2000 for detection of SARS-CoV-2. On days 8 and 9, the left caudal, right middle and right caudal lung lobes were evaluated, whereas on days 14 and 15, the left cranial, right cranial, and right middle lung lobes were evaluated. Tissue sections were analyzed by a blinded board-certified veterinary pathologist using an Olympus BX51 light microscope. Photomicrographs were taken on an Olympus DP73 camera.

#### Epitope mapping

Serum antibody epitope mapping competition assays were performed as previously described.^39,83^ Briefly, primary amine coupling was used to immobilize anti-histidine antibody on a Series S Sensor Chip CM5 (Cytiva) via His capture kit (Cytiva). His-tagged SARS-CoV-2 S-2P was then captured on the sensor surface. Following this, NHP sera (diluted 1:50) was flowed over both active and reference sensor surfaces. Active and reference sensor surfaces were regenerated between each analysis cycle. Human IgG monoclonal antibodies (mAb) used for these analyses include: S2-specific mAbs: S652-112 and S2P6, NTD-specific mAbs: 4-8, S652-118, N3C, and 5-7, RBD-specific mAbs B1-182, CB6, A20-29.1, A19-46.1, LY-COV555, A19-61.1, S309, A23-97.1, A19-30.1, A23-80.1, and CR3022, and SD1-specific mAb A19-36.1. Negative control antibody or competitor mAb was injected over both active and reference surfaces.

For analysis, sensorgrams were aligned to Y (Response Units) = 0, using Biacore 8K Insights Evaluation Software (Cytiva) beginning at the serum association phase. Relative “analyte binding late” report points (RU) were collected and used to calculate percent competition (% C) using the following formula: % C = [1 – (100 ^∗^ ((RU in presence of competitor mAb) / (RU in presence of negative control mAb)))]. Results are reported as percent competition.

### Quantification and statistical analysis

Comparisons between time points or within the same time point within a group are based on unpaired Student’s *t*-tests. Comparisons between groups within the same time points are based on one-way ANOVA with Tukey’s post-hoc test. Binding, neutralizing, and viral assays are log-transformed as appropriate and reported with medians and corresponding interquartile ranges where indicated. Adjustments for multiple comparisons were made when appropriate. All analyses are conducted using GraphPad Prism version 9.5.0 unless otherwise specified. The *p* values are shown in the figure as symbols defined in the figure legends, and the sample n is listed in corresponding figure legends. For all data presented, n=4 for individual boost cohorts and n=4-8 for controls and vaccinated NHP at pre-boost time points. If applicable, ‘ns’ denotes that the indicated comparison was not significant, with *p* > 0.05.

## Acknowledgments

This project was funded by the Intramural Research Program of the Vaccine Research Center (VRC), National Institutes of Allergy and Infectious Disease (NIAID), National Institutes of Health (NIH). We are grateful to PARI Pharma GmbH for providing the eFlow nebulizer for use in this study. We thank the VRC Production Program (VPP) for providing the WA-1 S-2P protein. VPP contributors include C. Anderson, V. Bhagat, J. Burd, J. Cai, K. Carlton, W. Chuenchor, N. Clbelli, G. Dobrescu, M. Figur, J. Gall, H. Geng, D. Gowetski, K. Gulla, L. Hogan, V. Ivleva, S. Khayat, P. Lei, Y. Li, I. Loukinov, M. Mai, S. Nugent, M. Pratt, E. Reilly, E. Rosales-Zavala, E. Scheideman, A. Shaddeau, A. Thomas, S. Upadhyay, K. Vickery, A. Vinitsky, C. Wang, C. Webber and Y. Yang. We thank Jay Noor from the Translational Research Program at the VRC for assistance with complete blood count tests. We also thank Serge Zouantcha and Elyse Teow for animal care and assistance with procedures and Brandon Narvaez for laboratory support at Bioqual. We thank Robert Coffman, Robert Janssen, Riccardo Manetti and Darren Campbell from Dynavax Technologies for providing CpG 1018 for use in this study.

## Author contributions

J.I.M, N.J.S., and R.A.S. designed experiments. J.I.M, S.F.A., B.J.F., D.A.W., K.E.F., M.G., D.R.F., E.L., S.P., J.M., A.M., C.G.L., C.C.H., M.R.B., L.M., A.R.H.., S.G., M.E.D.G., M.M., K.W.B., B.M.N., J.P.M.T., E.M., A.D., K.K., A.C., L.P., A.V.R., D.V., S.Y., Y.L., M.G.L., H.A., D.A.A., R.W., M.C.N., D.C.D., M.R., N.J.S., and R.A.S. performed, analyzed, and/or supervised experiments. J.I.M., K.E.F., S.F.A., and D.A.W. designed figures. A.C.M.B., M.S.S., A.L., A.V., S.Co., A.F., A.R., and S.Ca. provided critical reagents. J.I.M. wrote original draft of the manuscript. N.J.S., R.A.S., D.C.D., M.G., D.A.W. assisted with writing and provided feedback. All authors edited the manuscript and provided feedback on research.

---

**Figure S1.**
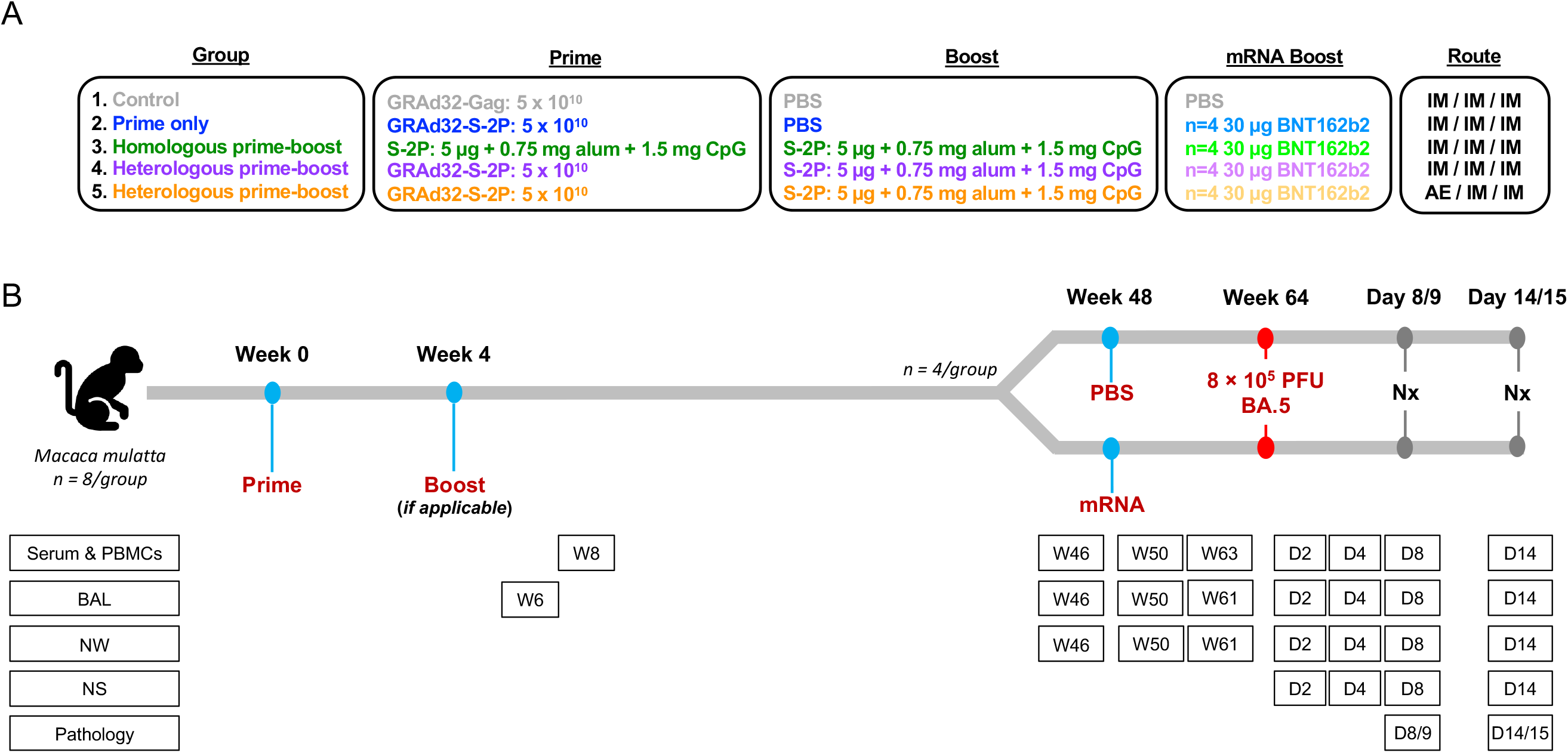
Experimental groups, timeline and sampling schedule, related to Figure 1. (A) Eight NHP per group were immunized with a variety of different SARS-CoV-2 immunogens. Group one was immunized with 5×10^10^ GRAd32-Gag. This group served as the control. Group two was immunized with 5×10^10^ GRAd32-S-2P. Group three was immunized with 5 μg adjuvanted S-2P (750 μg alum and 1500 μg CpG 1018; the same adjuvants were used when applicable). Group four was immunized with 5×10^10^ GRAd32-S-2P and with 5 μg adjuvanted S-2P. Group five was immunized with 5×10^10^ GRAd32-S-2P delivered via aerosol (AE) and with 5 μg adjuvanted S-2P. Phosphate buffered saline (PBS) was used as the placebo control as listed. (B) NHP were primed with the selected immunogen at week 0, and boosted, if applicable, at week 4. At week 48 (44 weeks after the second immunization or 48 weeks after prime only), the eight macaques in each group were subdivided into two groups of 4 (except control group) and immunized with 30 μg BNT162b2 (mRNA encoding S-2P). The week 0 and week 4 immunizations were delivered intramuscularly (IM) in 1 mL diluted in PBS (except aerosol GRAd32-S-2P) into the right deltoid, while the week 48 immunizations were delivered intramuscularly in 1 mL diluted in PBS into the right quadricep. At week 64, all NHP were challenged with 8 × 10^5^ PFU of BA.5. Samples were collected as listed. Abbreviations: BAL – Bronchoalveolar lavage, NW – nasal wash, NS – nasal swab, Nx – necropsy.

**Figure S2.**
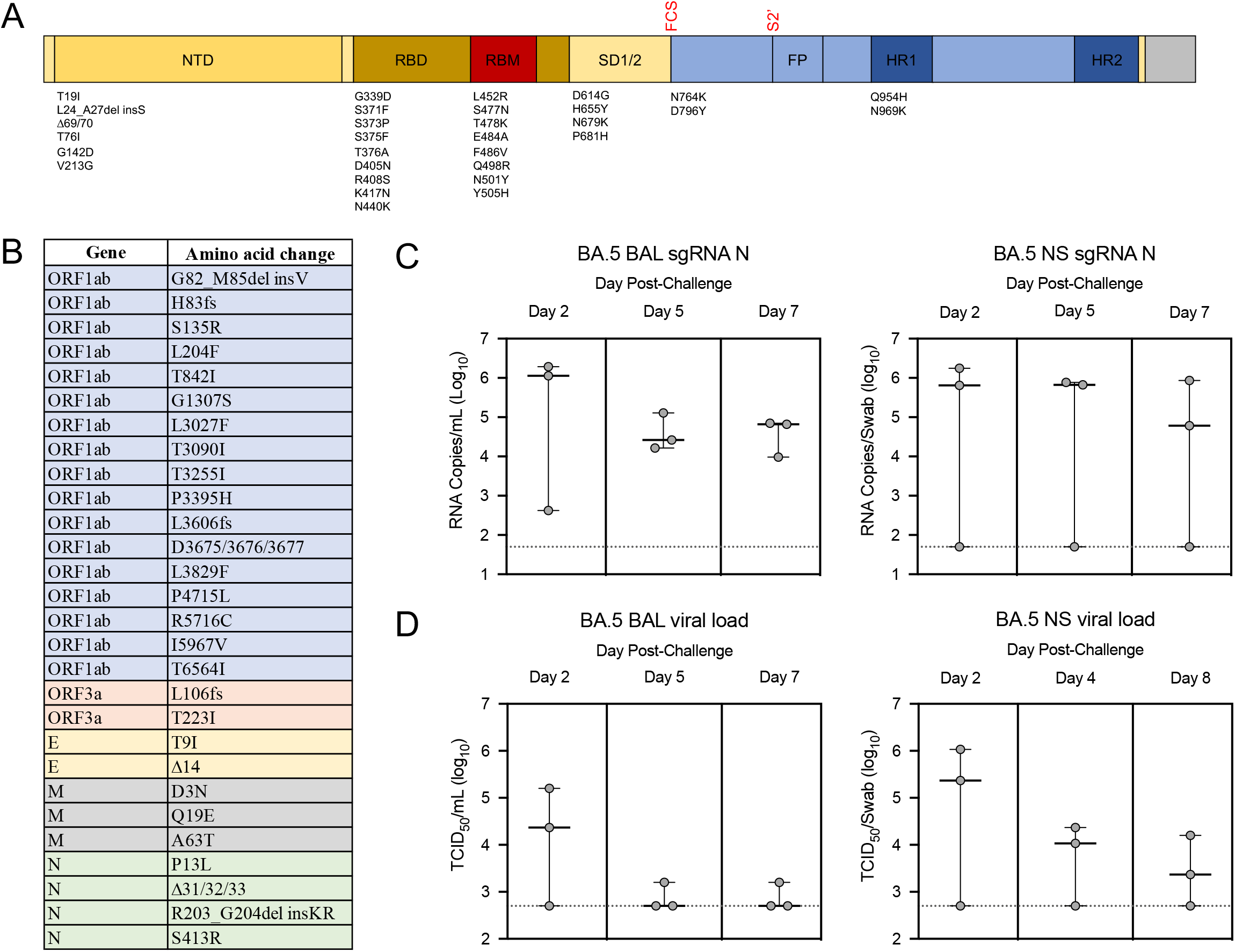
BA.5 challenge stock sequence and titration in NHP, related to Figure 1. (A and B) BA.5 stock was sequenced and aligned with Wuhan-Hu-1. (A) S gene only. Amino acid replacements listed above graphic. NTD, N-terminal domain; RBD, receptor binding domain; RBM, receptor binding motif; FP, fusion peptide; HR1, heptad repeat 1; HR2, heptad repeat 2; FCS, furin cleavage site; S2′, S2′ site. (B) Whole genome. (C–D) BA.5 stock was confirmed to be virulent in NHP. Circles indicate individual NHP. Error bars represent interquartile range with the median denoted by a dotted horizontal line. Assay limit of detection indicated by a horizontal dotted line. (C) BA.5 sgRNA_N copy numbers per mL of BAL or per swab in naïve NHP. (D) BA.5 TCID_50_ per mL of BAL or per swab in naïve NHP.

**Figure S3.**
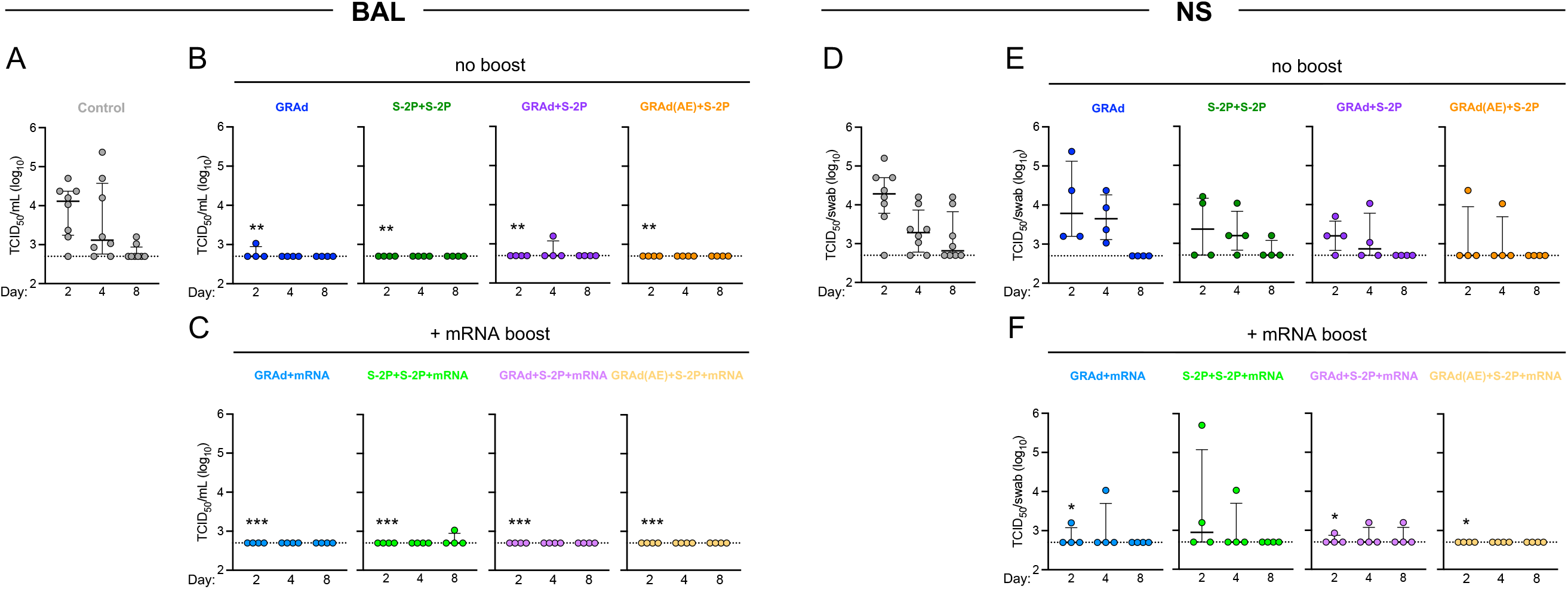
GRAd confers durable protection against BA.5 in the upper and lower airway, related to Figure 1 and 2. (A–F) BAL and NS was collected at days 2, 4 and 8 following challenge with 8 × 10^5^ PFU BA.5. (A) BA.5 TCID_50_ per mL in control NHP. (B) BA.5 TCID_50_ per mL in GRAd, S-2P+S-2P, GRAd+S-2P and GRAd(AE)-S-2P NHP. (C) BA.5 TCID_50_ per mL in GRAd, S-2P+S-2P, GRAd+S-2P and GRAd(AE)-S-2P NHP boosted with mRNA at week 48. (D) BA.5 TCID_50_ per swab in control NHP. (E) BA.5 TCID_50_ per swab in GRAd, S-2P+S-2P, GRAd+S-2P and GRAd(AE)-S-2P NHP. (F) BA.5 TCID_50_ per swab in GRAd, S-2P+S-2P, GRAd+S-2P and GRAd(AE)-S-2P NHP boosted with mRNA at week 48. Circles (A–F) indicate individual NHP. Error bars represent interquartile range with the median denoted by a horizontal line. Assay limit of detection indicated by a dotted horizontal line. Statistical analysis shown for corresponding timepoints between control and test group (e.g., ‘*’ symbols denote comparisons at day 2). * *p* <0.05, ** *p* <0.01, *** *p* <0.001. Eight control NHP and 4 immunized NHP per cohort.

**Figure S4.**
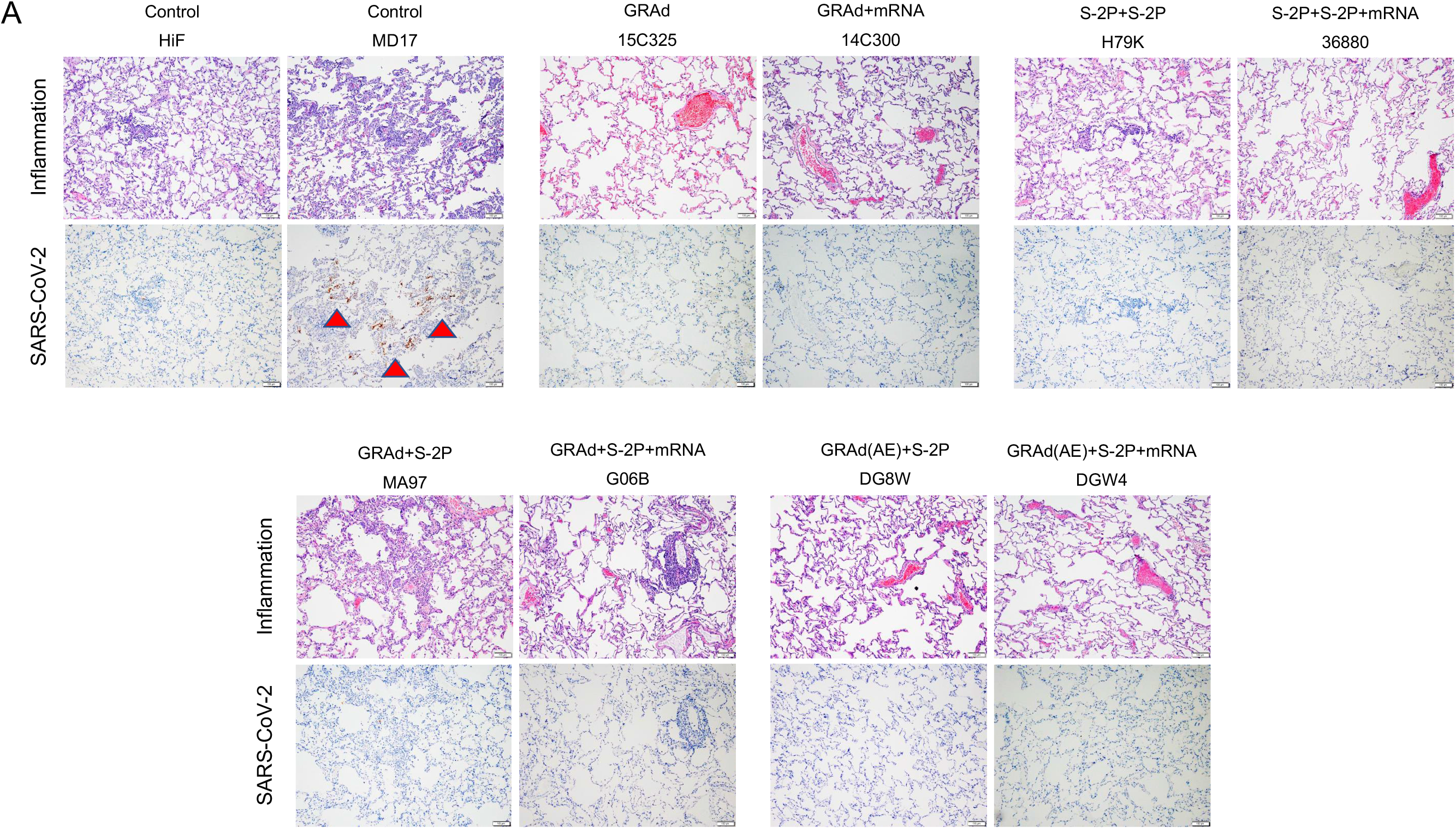
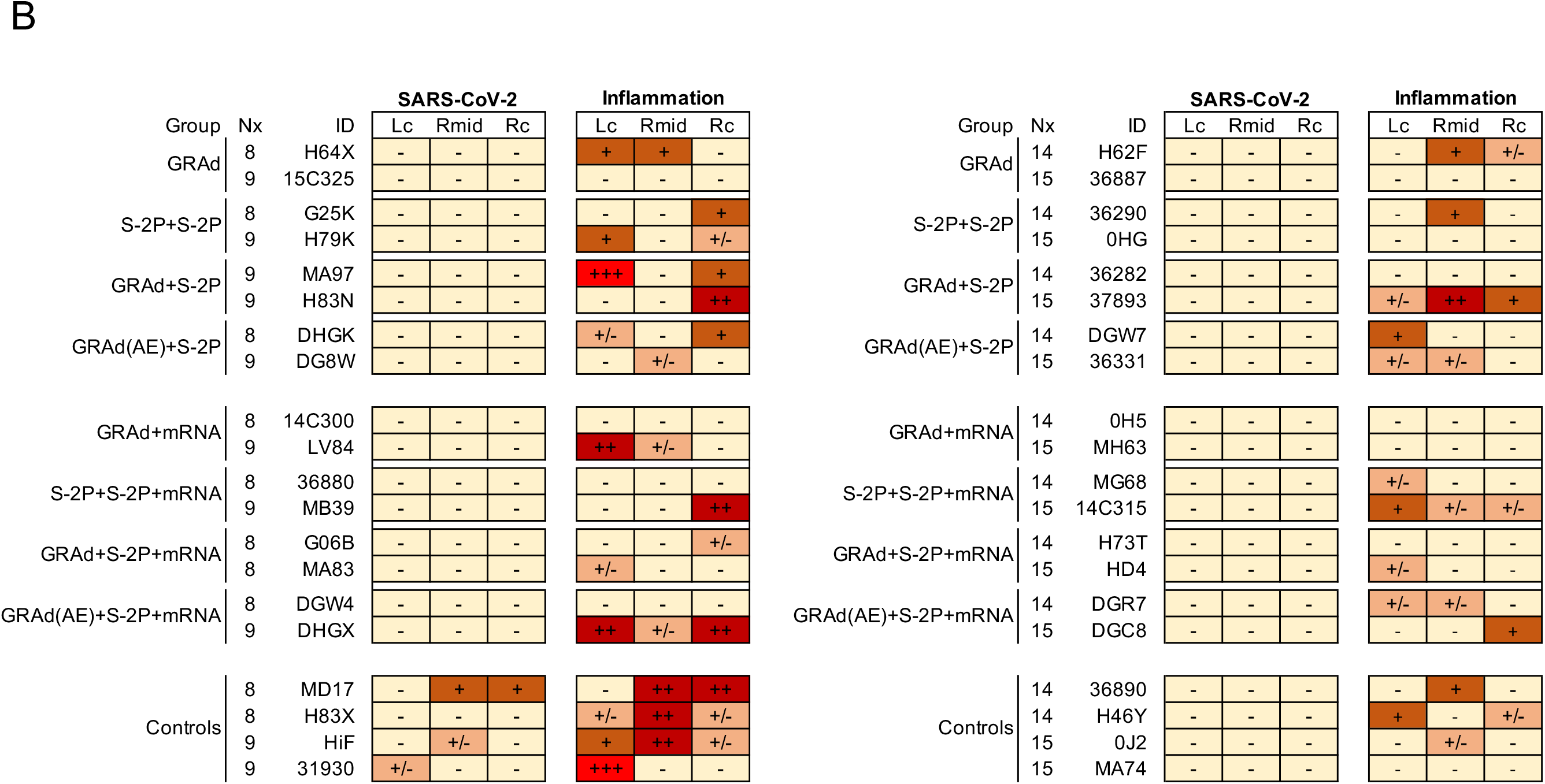
Viral antigen and pathology in the lung after challenge, related to Figure 1. (A and B) 2 NHP per group were euthanized on day 8 or 9 and 14 or 15 following challenge and tissue sections taken from the lung. (A) Top: Hematoxylin and eosin stain illustrating the extent of inflammation and cellular infiltrates. Bottom: Representative images indicating detection of SARS-CoV-2 N antigen by immunohistochemistry with a polyclonal anti-N antibody. Antigen-positive foci are marked by a red arrow. Images at 10x magnification with black bars for scale (100 μm). (B) SARS-CoV-2 antigen and inflammation scores in the left caudal lobe (Lc), right middle lobe (Rmid), and right caudal lobe (Rc) of the lung at days 8 or 9, and the left cranial lobe (Lc), right middle lobe (Rmid), and right cranial lobe (Rc) of the lung at days 14 or 15. Antigen scoring legend: − no antigen detected; +/− rare to occasional foci; + occasional to multiple foci; ++ multiple to numerous foci; +++ numerous foci. Inflammation scoring legend: − absent to minimal inflammation; +/− minimal to mild inflammation; + mild to moderate inflammation; ++ moderate-to-severe inflammation; +++ severe inflammation. Horizontal rows correspond to individual NHP.

**Figure S5.**
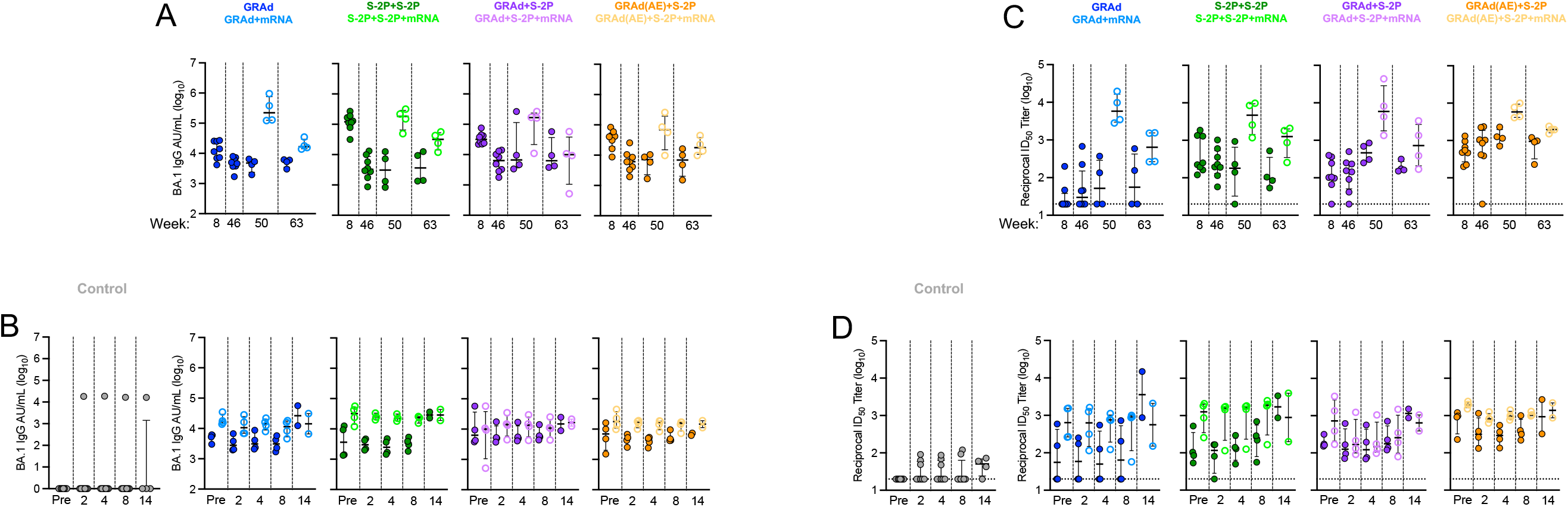
BA.1 binding and neutralization titers in serum pre-challenge and post-challenge, related to Figure 3. (A and C) Sera were collected at week 8, 46, 50 and 63. (B and D) Sera were collected at days 2, 4, 8 and 14 following challenge with BA.5. (A) IgG-binding titers to BA.1 S expressed in AU/mL pre-challenge. (B) IgG-binding titers to BA.1 S expressed in AU/mL post-challenge. (C) Neutralizing titers to BA.1 lentiviral pseudovirus expressed as the reciprocal ID_50_ pre-challenge. (D) Neutralizing titers to BA.1 lentiviral pseudovirus expressed as the reciprocal ID_50_ post-challenge. Circles (A–D) represent individual NHP. Error bars represent the interquartile range with the median denoted by a horizontal black line. Assay limit of detection indicated by a horizontal dotted line which may fall below the depicted range. Vertical dashed lines are for visualization purposes only. Eight immunized NHP, split into 2 cohorts of 4 NHP post mRNA boost. Eight control NHP (4 at day 14), 8 immunized NHP at week 8 and 46, 4 immunized NHP at week 50 and 63, 4 immunized NHP at days 2, 4 and 8, and 2 immunized NHP at day 14.

**Figure S6.**
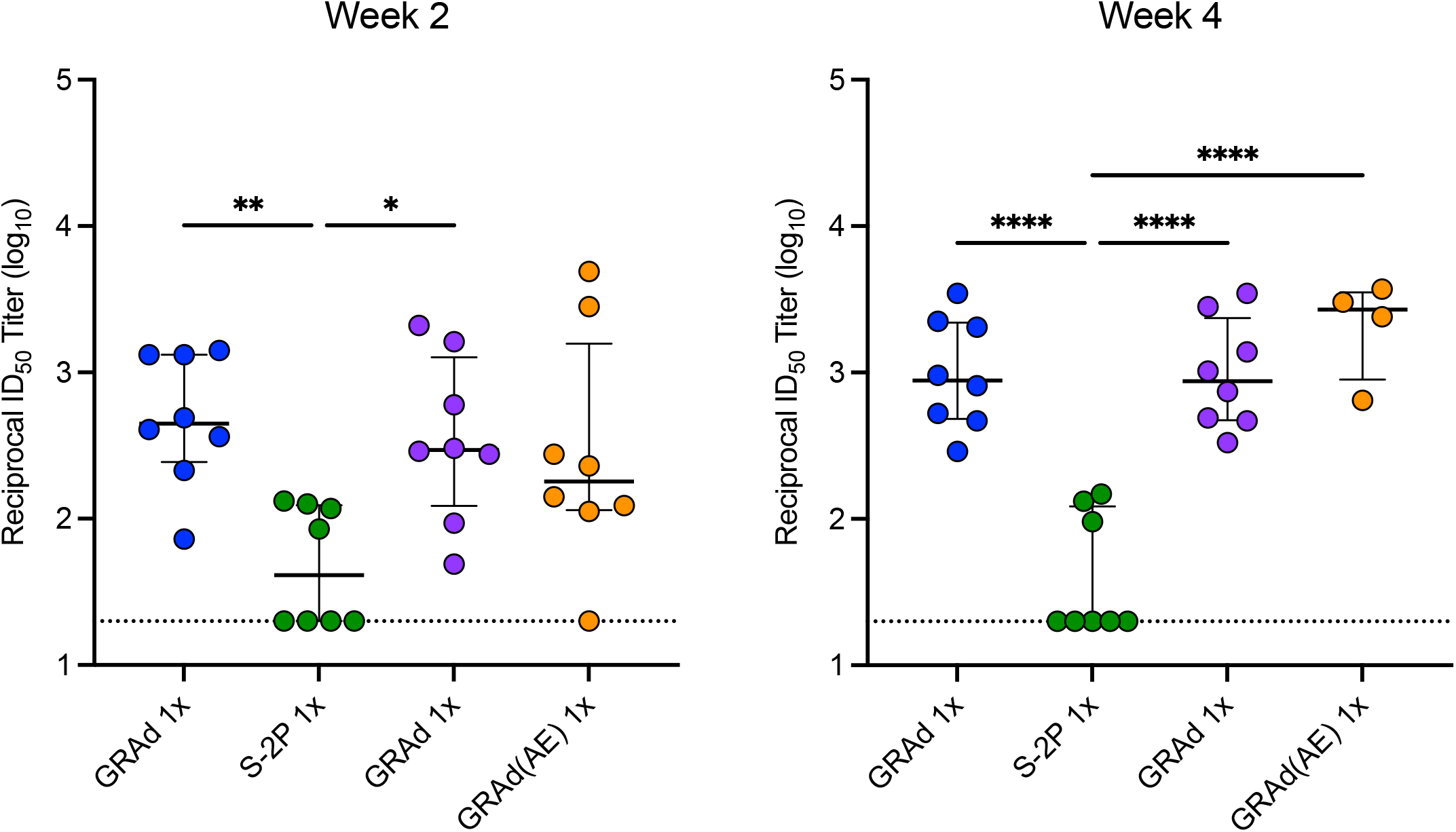
D614G neutralization titers in serum at week 2 and 4, related to Figure 3. Sera were collected at week 2 and 4. Neutralizing titers to ancestral D614G lentiviral pseudovirus expressed as the reciprocal ID_50_. Circles represent individual NHP. Error bars represent the interquartile range with the median denoted by a horizontal black line. Assay limit of detection indicated by a horizontal dotted line. Eight immunized NHP (except for week 4 GRAd(AE) 1x, only 4 out of the 8 NHP were not bled at this timepoint due to blood sampling limits). Statistical analysis shown for corresponding timepoint between groups. * *p* <0.05, ** *p* <0.01, **** *p* <0.0001.

**Figure S7.**
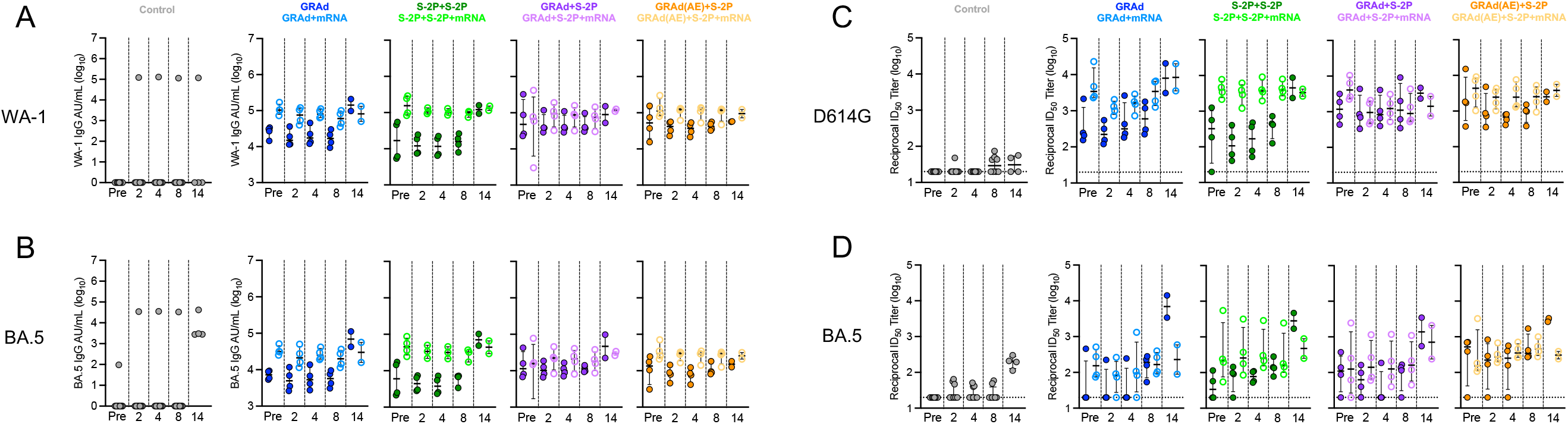
WA-1 or D614G and BA.5 binding and neutralization titers in serum following challenge, related to Figure 3. (A–D) Sera were collected at days 2, 4, 8 and 14 following challenge with BA.5. (A and B) IgG-binding titers to WA-1 and BA.5 S expressed in AU/mL post-challenge. (C and D) Neutralizing titers to WA-1 and BA.5 lentiviral pseudovirus expressed as the reciprocal ID_50_ post-challenge. Circles (A–D) represent individual NHP. Error bars represent the interquartile range with the median denoted by a horizontal black line. Assay limit of detection indicated by a horizontal dotted line which may fall below the depicted range. Vertical dashed lines are for visualization purposes only. Eight immunized NHP, split into 2 cohorts of 4 NHP post mRNA boost. Eight control NHP (4 at day 14), 8 immunized NHP at week 8 and 46, 4 immunized NHP at week 50 and 63, 4 immunized NHP at days 2, 4 and 8, and 2 immunized NHP at day 14.

**Figure S8.**
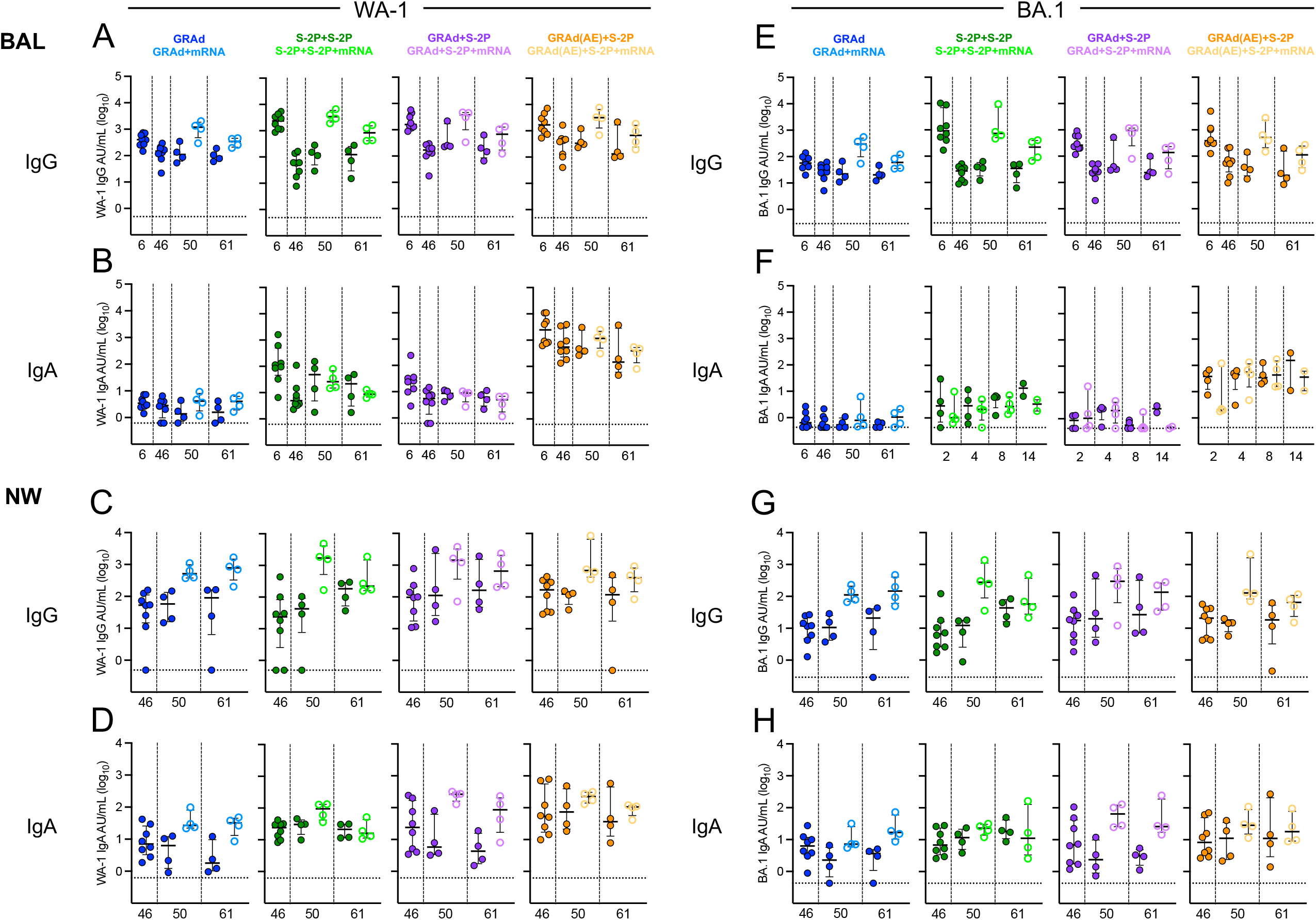
WA-1 and BA.1 IgG and IgA binding titers in BAL and NW prior to challenge, related to Figure 4. (A–H) BAL (A and B) was collected at week 6, 46, 50 and 61 and NW (C and D) was collected at week 46, 50, and 61. (A and B) IgG (A) and IgA (B) antibody binding titers to WA-1 expressed in AU/mL in BAL. (C and D) IgG (C) and IgA (D) antibody binding titers to WA-1 expressed in AU/mL in NW. (E and F) IgG (A) and IgA (B) antibody binding titers to BA.1 expressed in AU/mL in BAL. (G and H) IgG (C) and IgA (D) antibody binding titers to BA.1 expressed in AU/mL in NW. Circles (A–G) represent individual NHP. Error bars represent the interquartile range with the median denoted by a horizontal black line. Assay limit of detection indicated by a horizontal dotted. Vertical dashed lines are for visualization purposes only. Eight vaccinated NHP, split into 2 cohorts of 4 NHP post mRNA boost. Eight immunized NHP at week 6 and 46, 4 immunized NHP at week 50 and 61.

**Figure S9.**
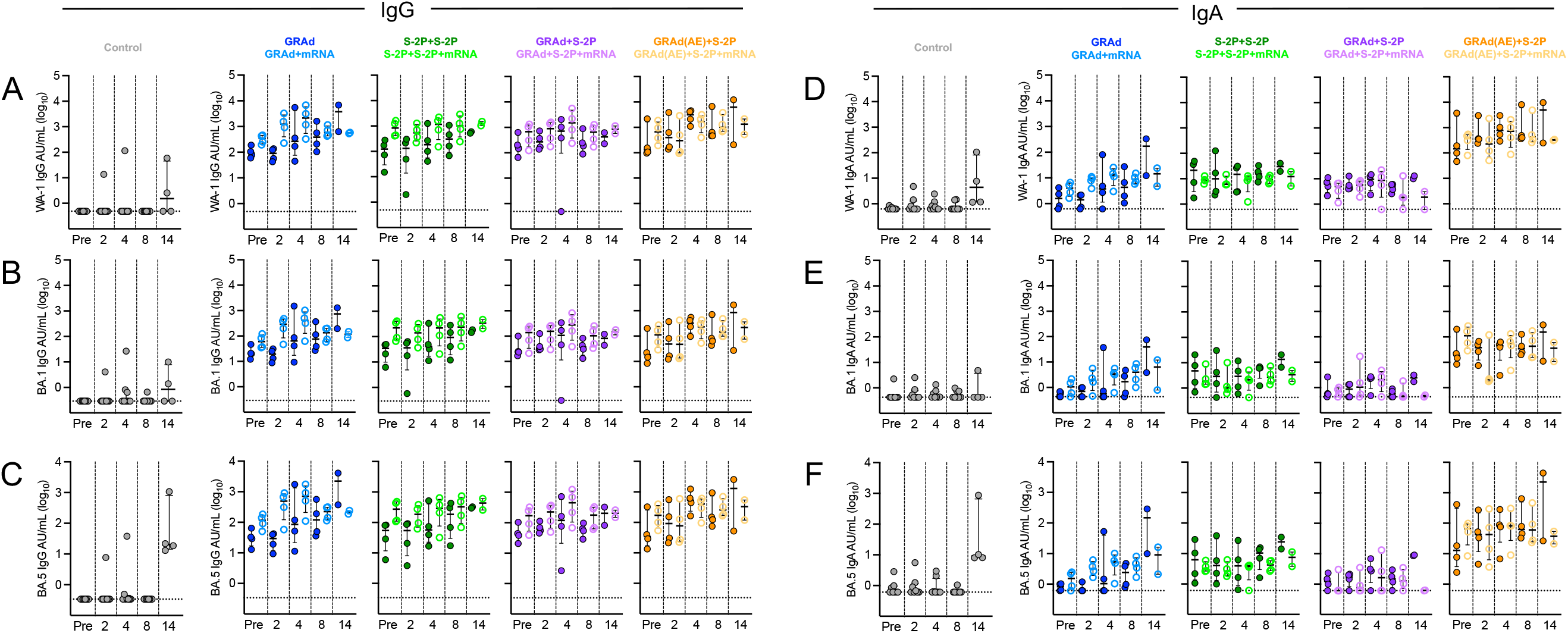
IgG and IgA binding titers in BAL following challenge, related to Figure 4. (A–F) BAL was collected at days 2, 4, 8 and 14 following challenge. (A–C) IgG antibody binding titers to WA-1 (A), BA.1 (B) and BA.5 (C) expressed in AU/mL in BAL. (D–F) IgA antibody binding titers to WA-1 (A), BA.1 (B) and BA.5 (C) expressed in AU/mL in BAL. Circles (A–F) represent individual NHP. Error bars represent the interquartile range with the median denoted by a horizontal black line. Assay limit of detection indicated by a horizontal dotted line. Vertical dashed lines are for visualization purposes only. Eight vaccinated NHP, split into 2 cohorts of 4 NHP post mRNA boost. Eight control NHP (4 at day 14), 4 immunized NHP at days 2, 4 and 8, 2 immunized NHP at day 14. For reference, the week 61 BAL IgG or IgA titer was included in the graphs as “pre”.

**Figure S10.**
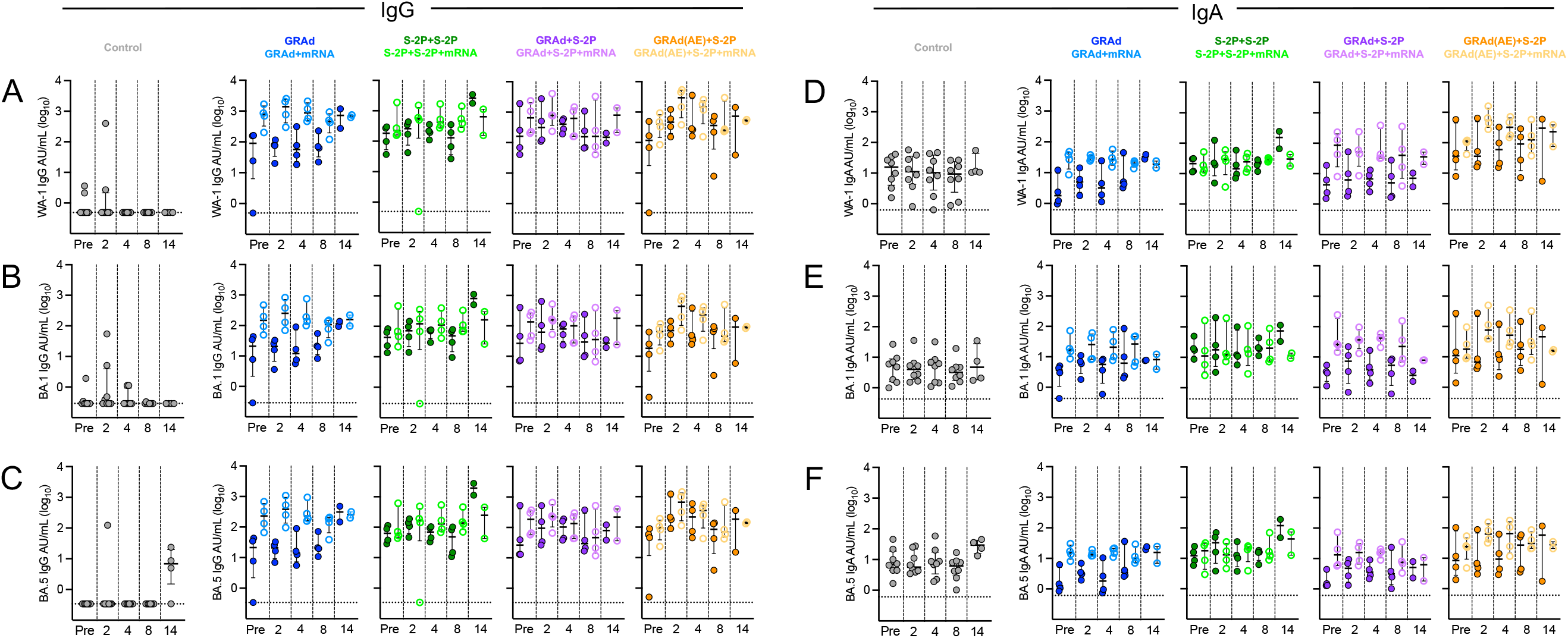
IgG and IgA binding titers in NW following challenge, related to Figure 4. (A–F) NW was collected at days 2, 4, 8 and 14 following challenge. (A–C) IgG antibody binding titers to WA-1 (A), BA.1 (B) and BA.5 (C) expressed in AU/mL in NW. (D–F) IgA antibody binding titers to WA-1 (A), BA.1 (B) and BA.5 (C) expressed in AU/mL in NW. Circles (A–F) represent individual NHP. Error bars represent the interquartile range with the median denoted by a horizontal black line. Assay limit of detection indicated by a horizontal dotted line. Vertical dashed lines are for visualization purposes only. Eight vaccinated NHP, split into 2 cohorts of 4 NHP post mRNA boost. Eight control NHP (4 at day 14), 4 immunized NHP at days 2, 4 and 8, 2 immunized NHP at day 14. For reference, the week 61 NW IgG or IgA titer was included in the graphs as “pre”.

**Figure S11.**
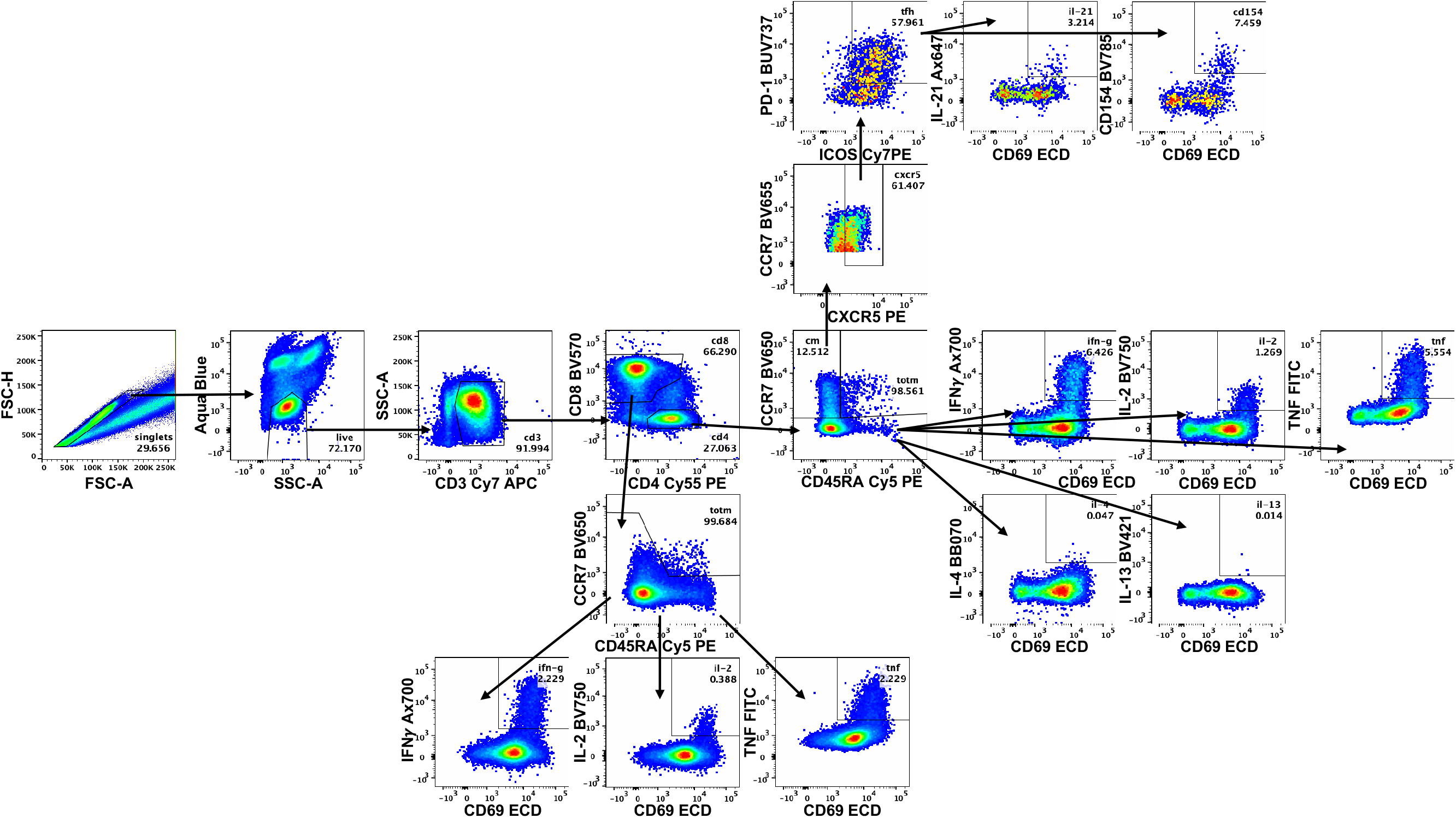
T cell gating strategy, related to Figure 5. Representative flow cytometry plots showing gating strategy for T cells in Figures 5, S11 and S12. Cells were gated as singlets and live cells on forward and side scatter and a live/dead aqua blue stain. CD3+ events were gated as CD4+ or CD8+ T cells. Total memory CD8+ T cells were selected based on expression of CCR7 and CD45RA, and SARS-CoV-2 S-specific memory CD8+ T cells were gated according to co-expression of CD69 and IL-2, TNF or IFNγ. The CD4+ events were defined as naive, total memory, or central memory according to expression of CCR7 and CD45RA, and CD4+ cells with a T_h_1 phenotype were defined as memory cells that co-expressed CD69 and IL-2, TNF or IFNγ. CD4+ cells with a T_h_2 phenotype were defined as memory cells that co-expressed CD69 and IL-4 or IL-13. T_fh_ cells were defined as central memory CD4+ T cells that expressed CXCR5, ICOS, and PD-1. T_fh_ cells were further characterized as IL-21+ and CD69+ or CD40L+ (CD154) and CD69+.

**Figure S12.**
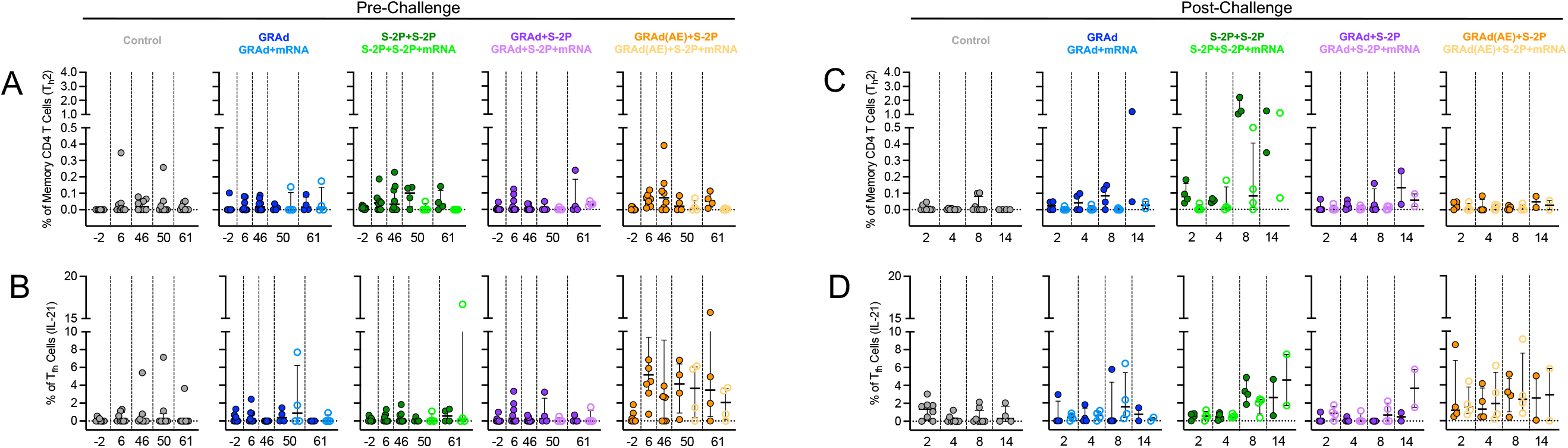
T_h_2 and IL-21+ T_fh_ cells in BAL, related to Figure 5. (A–B) BAL was collected at week −2, 6, 46, 50 and 61. Cells were stimulated with SARS-CoV-2 S1 and S2 peptide pools (WA1) and then measured by intracellular cytokine staining. (C–D) BAL was collected at days 2, 4, 8, 50 and 14 following challenge. Cells were stimulated with SARS-CoV-2 S1 and S2 peptide pools (WA-1) and then measured by intracellular cytokine staining. (A, C) Percentage of memory CD4+ T cells with T_h_2 markers (IL-4 or IL-13) following stimulation. (B, D) Percentage of T_fh_ cells that express IL-21. Circles in (A–D) indicate individual NHP. Error bars represent the interquartile range with the median denoted by a horizontal black line. Dotted lines set at 0%. Reported percentages may be negative due to background subtraction and may extend below the range of the y axis. Eight vaccinated NHP, split into 2 cohorts of 4 NHP post mRNA boost. Eight control NHP at days 2, 4 and 8, four at day 14.

**Figure S13.**
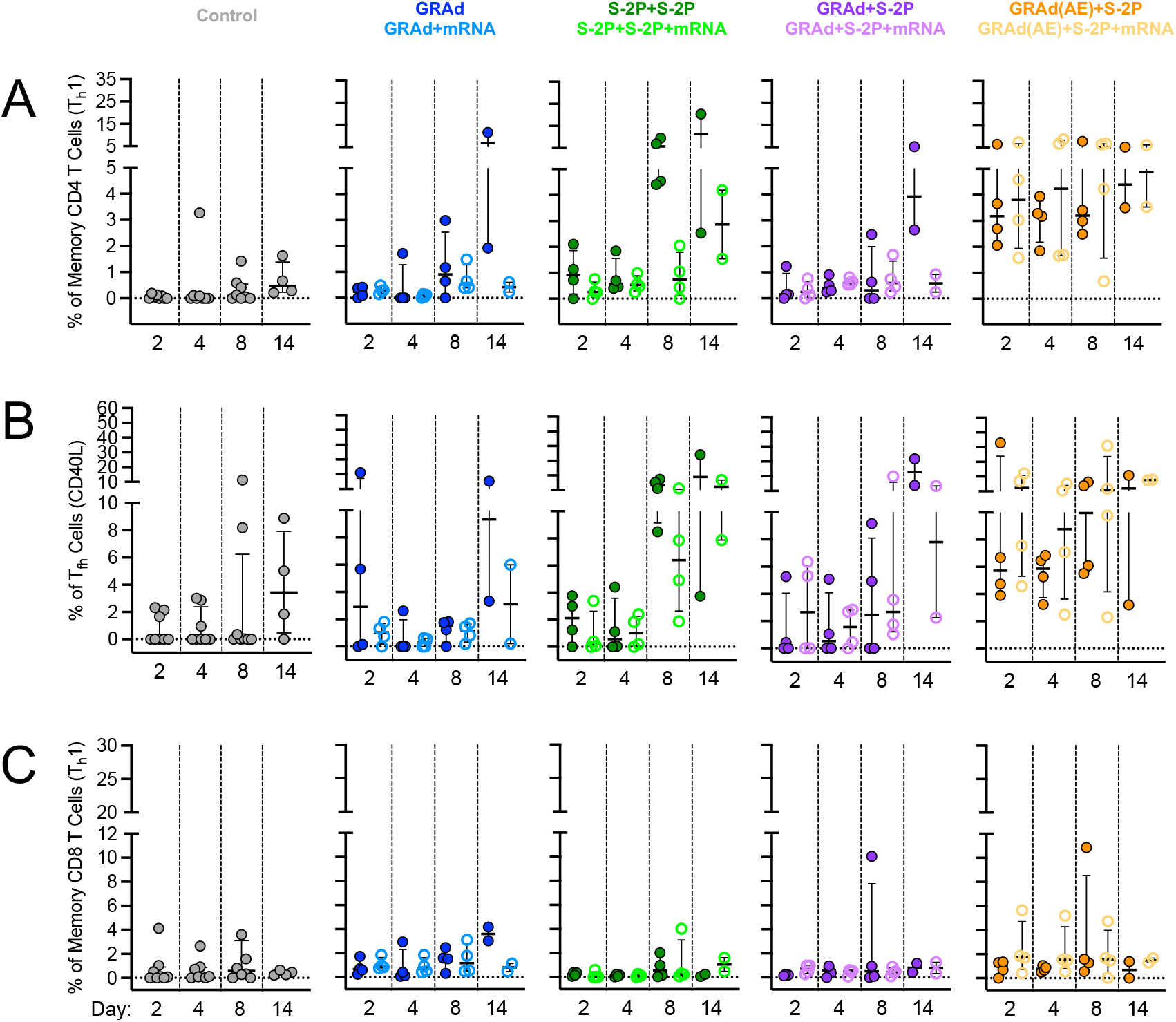
T cell in BAL following challenge, related to Figure 5. (A–C) BAL was collected at days 2, 4, 8 and 14 following challenge. Cells were stimulated with SARS-CoV-2 S1 and S2 peptide pools (WA-1) and then measured by intracellular cytokine staining. (A) Percentage of memory CD4+ T cells with Th1 markers (IL-2, TNF, or IFNγ). (B) Percentage of T_fh_ cells that express CD40L. (C) Percentage of CD8 T cells expressing IL-2, TNF, or IFNγ. Circles in (A–D) indicate individual NHP. Error bars represent the interquartile range with the median denoted by a horizontal black line. Dotted lines set at 0%. Reported percentages may be negative due to background subtraction and may extend below the range of the y axis. Eight vaccinated NHP, split into 2 cohorts of 4 NHP post mRNA boost. Eight control NHP (4 at day 14), 4 immunized NHP at days 2, 4 and 8, 2 immunized NHP at day 14.

**Figure S14.**
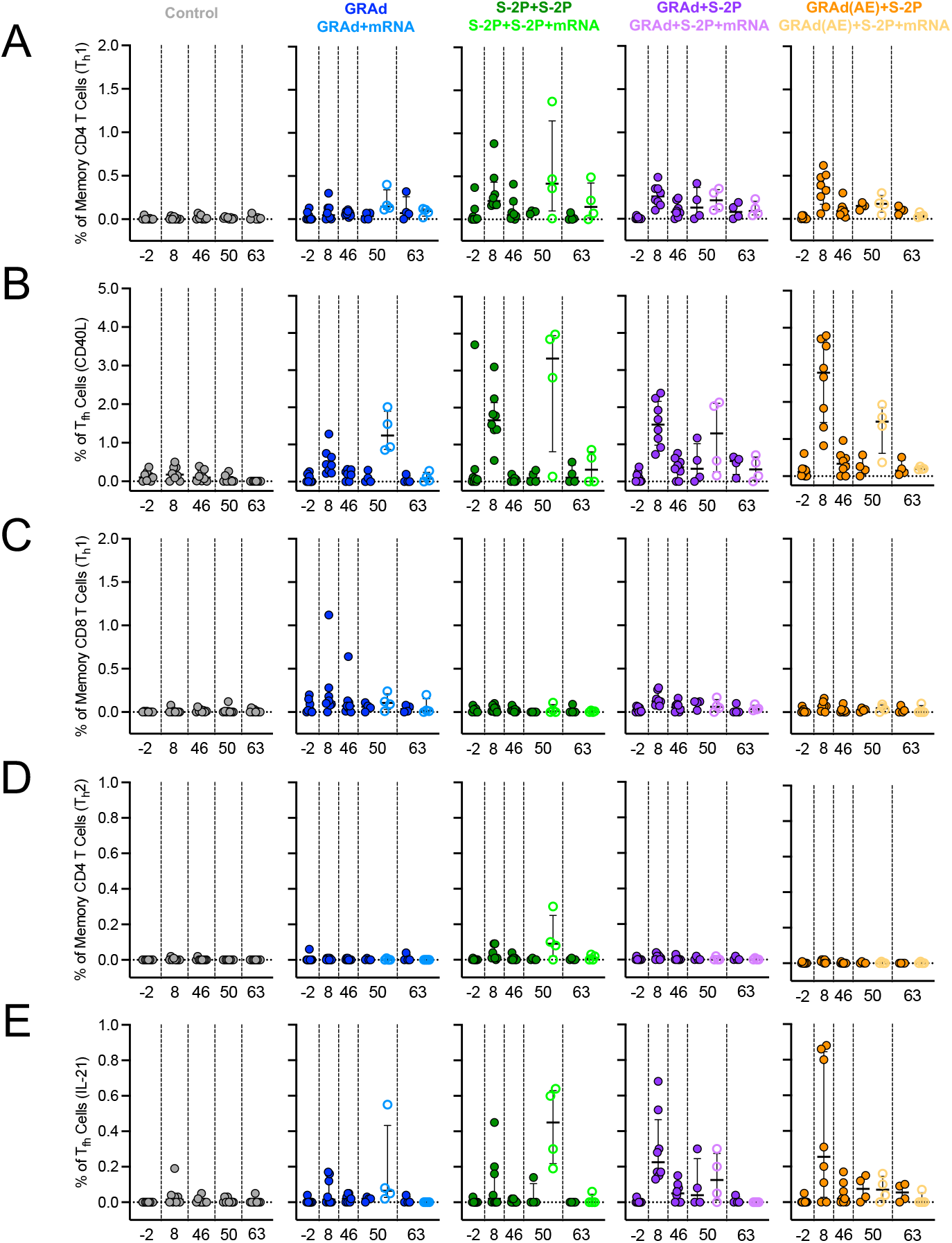
T cell in blood prior to challenge, related to Figure 5. (A–E) Peripheral blood mononuclear cells (PBMCs) were collected at week −2, 8, 46, 50 and 63. Cells were stimulated with SARS-CoV-2 S1 and S2 peptide pools (WA-1) and then measured by intracellular cytokine staining. (A) Percentage of memory CD4+ T cells with Th1 markers (IL-2, TNF, or IFNγ). (B) Percentage of T_fh_ cells that express CD40L. (C) Percentage of CD8 T cells expressing IL-2, TNF, or IFNγ. (D) Percentage of memory CD4+ T cells with T_h_2 markers (IL-4 or IL-13). (E) Percentage of T_fh_ cells that express IL-21. Circles in (A–E) indicate individual NHP. Error bars represent the interquartile range with the median denoted by a horizontal black line. Dotted lines set at 0%. Reported percentages may be negative due to background subtraction and may extend below the range of the y axis. Eight vaccinated NHP, split into 2 cohorts of 4 NHP post mRNA boost. Eight control NHP at days 2, 4 and 8, four at day 14.

**Figure S15.**
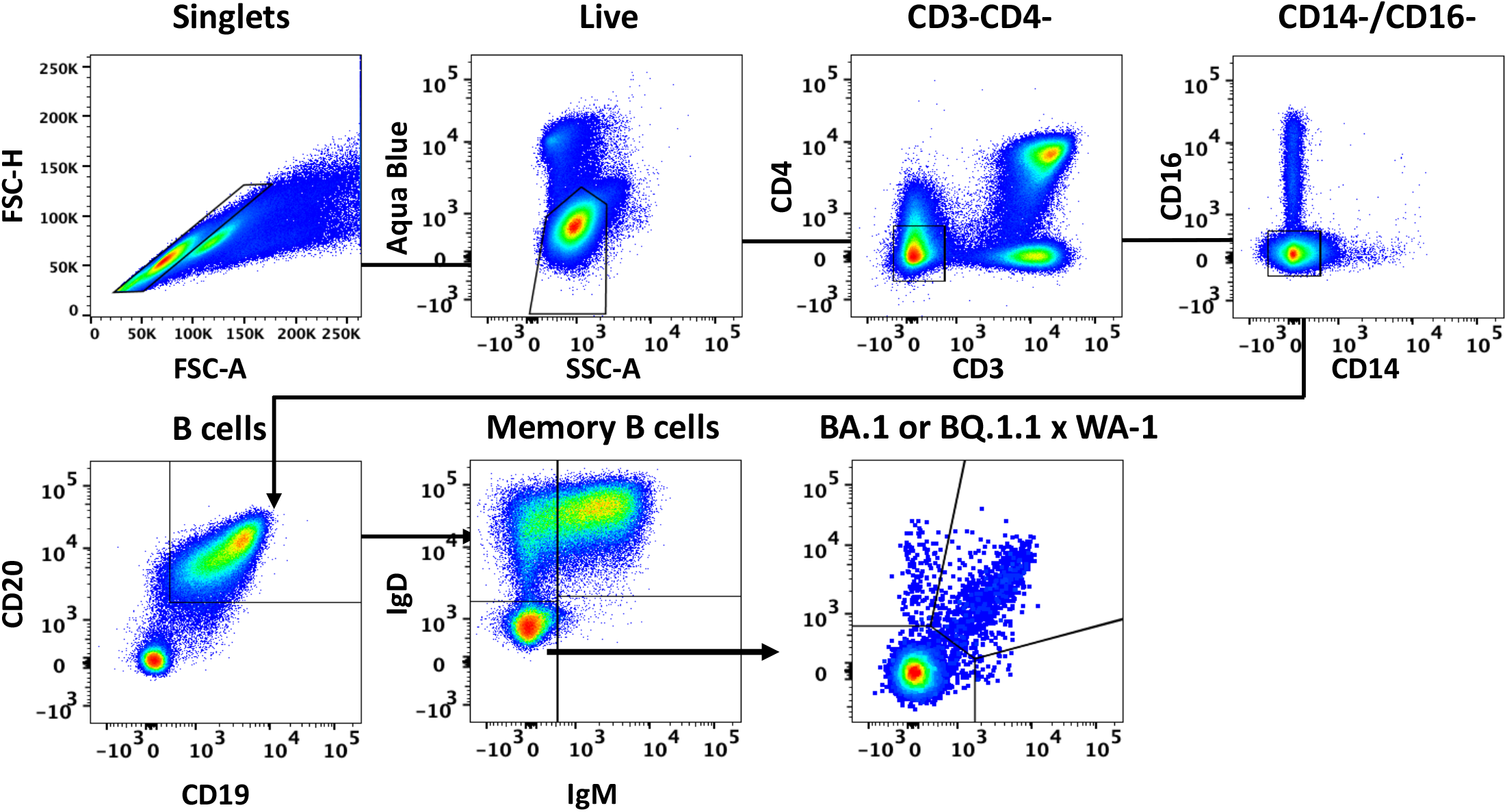
B cell gating strategy, related to Figure 6. Representative flow cytometry plots showing gating strategy for B cells in Figures 6 and S16. Cells were gated as singlets and live cells on forward and side scatter and a live/dead aqua blue stain. CD3-, CD4-cells were then gated on absence of CD14 and CD16 expression and positive expression of CD20 and CD19. Memory B cells were selected based on lack of IgD or IgM. Finally, memory B cells of variant S-2P (WA-1 and BA.5 or WA-1 and BQ.1.1) probes were used to determine binding specificity.

**Figure S16:**
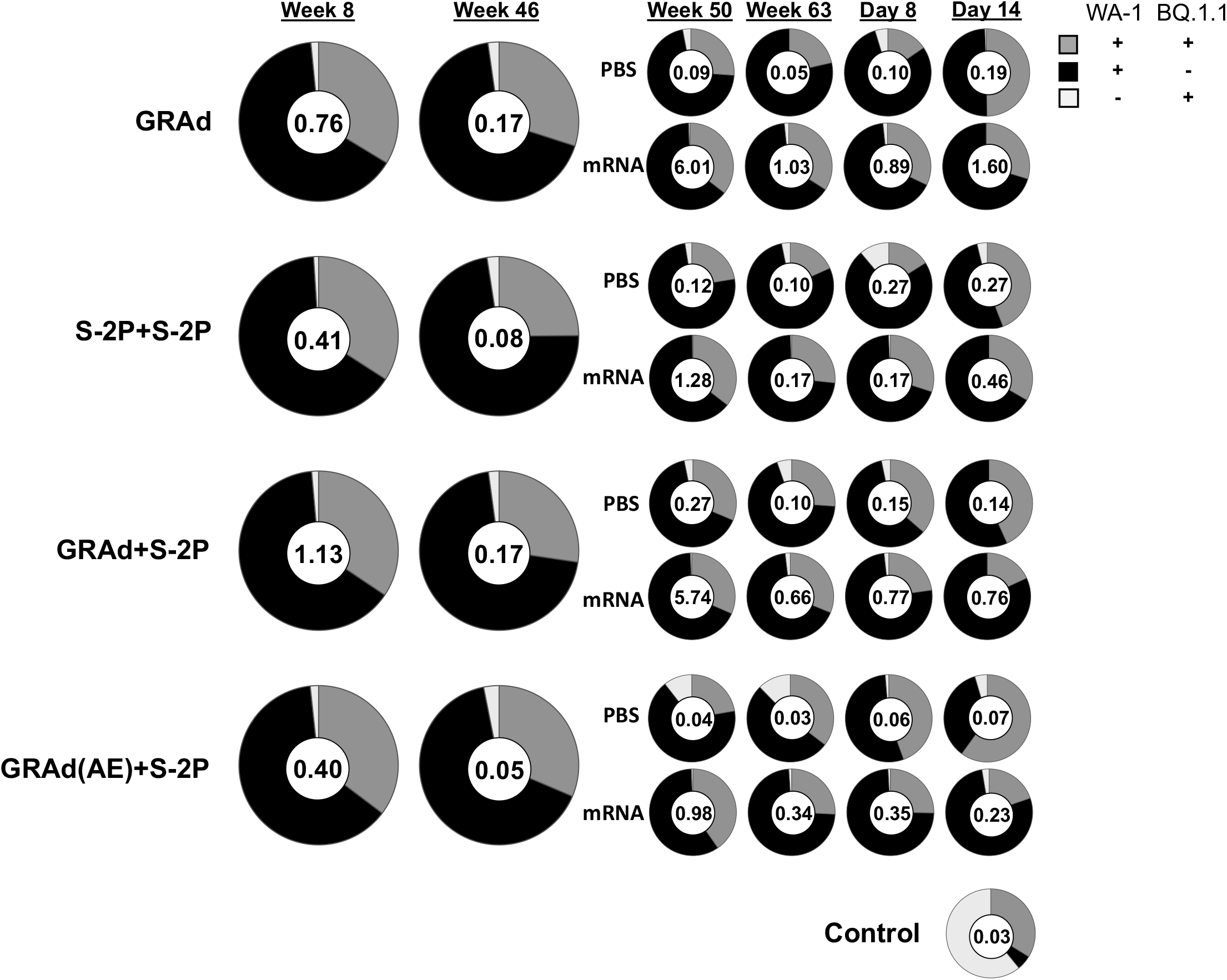
WA-1 and BQ.1.1 cross-reactive S-2P specific memory B cells following immunization, related to Figure 6. Pie charts indicate the frequency (numbered circle at the center) and proportion of total S-binding memory B cells that are dual specific for WA-1 and BQ.1.1 (dark gray), specific for WA-1 (black), or specific for BQ.1.1 (light gray) for all NHP in each group and timepoint at week 8, 46, 50 and 63 post-immunization, and days 8 and 14 post-challenge. Seven or eight NHP per group at week 8 and 46, 3-4 NHP per group at week 50 and 63, and day 8, 1-2 NHP at day 14.

